# piRNA pathway is essential for generating functional oocytes in golden hamster

**DOI:** 10.1101/2021.03.21.434510

**Authors:** Hongdao Zhang, Fengjuan Zhang, Jinghua Chen, Mingzhi Li, Xiaolong Lv, Yali Xiao, Zhaozhen Zhang, Li Hou, Yana Lai, Wen Xiao, Aihua Zhang, Heling Fu, Jianli Zhou, Feiyang Diao, Aimin Shi, Youqiang Su, Wentao Zeng, Ligang Wu, Jianmin Li

**Affiliations:** State Key Laboratory of Molecular Biology, Shanghai Key Laboratory of Molecular Andrology, Center for Excellence in Molecular Cell Science, Shanghai Institute of Biochemistry and Cell Biology, Chinese Academy of Sciences, University of Chinese Academy of Sciences, Shanghai 200031, China; Model Animal Research Center of State Key Laboratory of Reproductive Medicine, Jiangsu Laboratory Animal Center, Jiangsu Animal Experimental Center of Medicine and Pharmacy, Jiangsu Animal of Model, Department of Cell Biology, Animal Core facility, Key Laboratory of Model Animal, Collaborative Innovation Center for Cardiovascular Disease Translational Medicine, Nanjing Medical University, Nanjing 211166, China; Clinical Center of Reproductive Medicine, the First Hospital Affiliated with Nanjing Medical University, Nanjing, China

## Abstract

Piwi-interacting RNAs (piRNAs) are small RNAs predominantly expressed in germ cells that are critical for gametogenesis in various species. However, PIWI-deficient female mice are fertile and mouse oocytes express a panel of small RNAs that do not appear widely representative of mammals, and piRNA function in the oogenesis of other mammals has therefore remained elusive. Recent studies revealed the small RNA and *PIWI* transcriptional profiles in golden hamster oocytes more closely resemble that of humans than mice. Herein, we generated *PIWIL1*-, *PLD6-* and *MOV10L1-*deficient golden hamsters and found that all female mutants were sterile, with embryos arrested at the two-cell stage. In *PIWIL1* mutant oocytes, we observed transposon accumulation and broad transcriptomic dysregulation, while zygotic gene activation was impaired in early embryos. Intriguingly, PIWIL1-piRNAs exhibited a unique, preferential silencing of endogenous retroviruses (ERVs), whereas silencing LINE1s depended on both PIWIL1- and PIWIL3-piRNAs. Moreover, we showed that piRNAs participate in the degradation of maternal mRNAs in MII oocytes and embryos via partially complementary targets. Together, our findings demonstrate that piRNAs are indispensable for generating functional oocytes in golden hamster, and show the informative value of this model for functional and mechanistic investigations of piRNAs, especially those related to female infertility.

## Introduction

Over the past two decades, substantial progress has been made towards understanding the transcriptional and post-transcriptional regulatory systems in gene expression mediated by small RNAs. Among these, PIWI-interacting RNAs (piRNAs) are single-stranded small RNAs, 18-30 nt in length, that are predominantly expressed in germ cells and bind with PIWI proteins to form piRNA-induced silencing complexes (piRISCs), which provide diverse and indispensable functions in gametogenesis (*1-6*). The primary function of the piRNA pathway is to suppress active transposable elements (TEs) by post-transcriptional gene silencing (PTGS) through piRISC-mediated slicer activity, as well as transcriptional gene silencing (TGS) through guiding DNA methylation or histone modification (*7, 8*). Moreover, increasing evidence has shown that the piRNA pathway could also regulate mRNAs and lncRNAs (*9-12*), and are functional in tissue regeneration (*13, 14*), tumor biology (*15, 16*), and embryogenesis (*17-19*).

In *Drosophila* and *Zebrafish*, disruption of any of the PIWI paralogs severely impairs the fertility of both males and females (*20-23*). In mammals, the piRNA pathway has been primarily studied in mice, whose genome encodes three *PIWIs*, including *PIWIL1* (*MIWI*), *PIWIL2* (*MILI*), and *PIWIL4* (*MIWI2*). Disruption of any *PIWIs* in mice leads to spermatogenesis arrest and sterility in males *(24-26)*. However, female mice remain fertile in the absence of *PIWI*s, as well as other genes that are critical for piRNA biogenesis (*24-30*). Unlike many other mammals, endo-siRNAs are the most abundant small RNAs in mouse oocytes, which might be attributable to the expression of a unique isoform of Dicer (Dicer^O^) that is highly effective in producing endo-siRNAs, while the abundance of piRNAs is dwarfed in contrast (*31-33*). These differences suggest that the dispensability of the piRNA pathway in female fertility previously reported in mice may not be representative of all mammals, including humans, and that the role of piRNAs warrants closer examination in other animals. Golden hamster (*Mesocricetus auratus*) belongs to the *Cricetidae* family of rodents and has been used as a model in studying development and reproductive biology for several decades (*34*). Moreover, the golden hamster genome encodes all four *PIWIs*, with an oocyte small RNA transcriptional profile that more closely resembles the human profile than the mouse (Fig. 1A-B) (*1, 35*). In particular, piRNAs and PIWIs are highly expressed, thus providing an informative animal model to investigate the potential involvement of the piRNA pathway in female gametogenesis. In this study, we demonstrate the essential function of the piRNA pathway in female reproduction in golden hamster. Furthermore, we provide evidence that PIWIL1 and piRNAs are functional in mediating transposon silencing and degradation of maternal mRNAs, and that they play an essential role in oogenesis and the initiation of ZGA to sustain the development of early embryos.

**Fig. 1.**
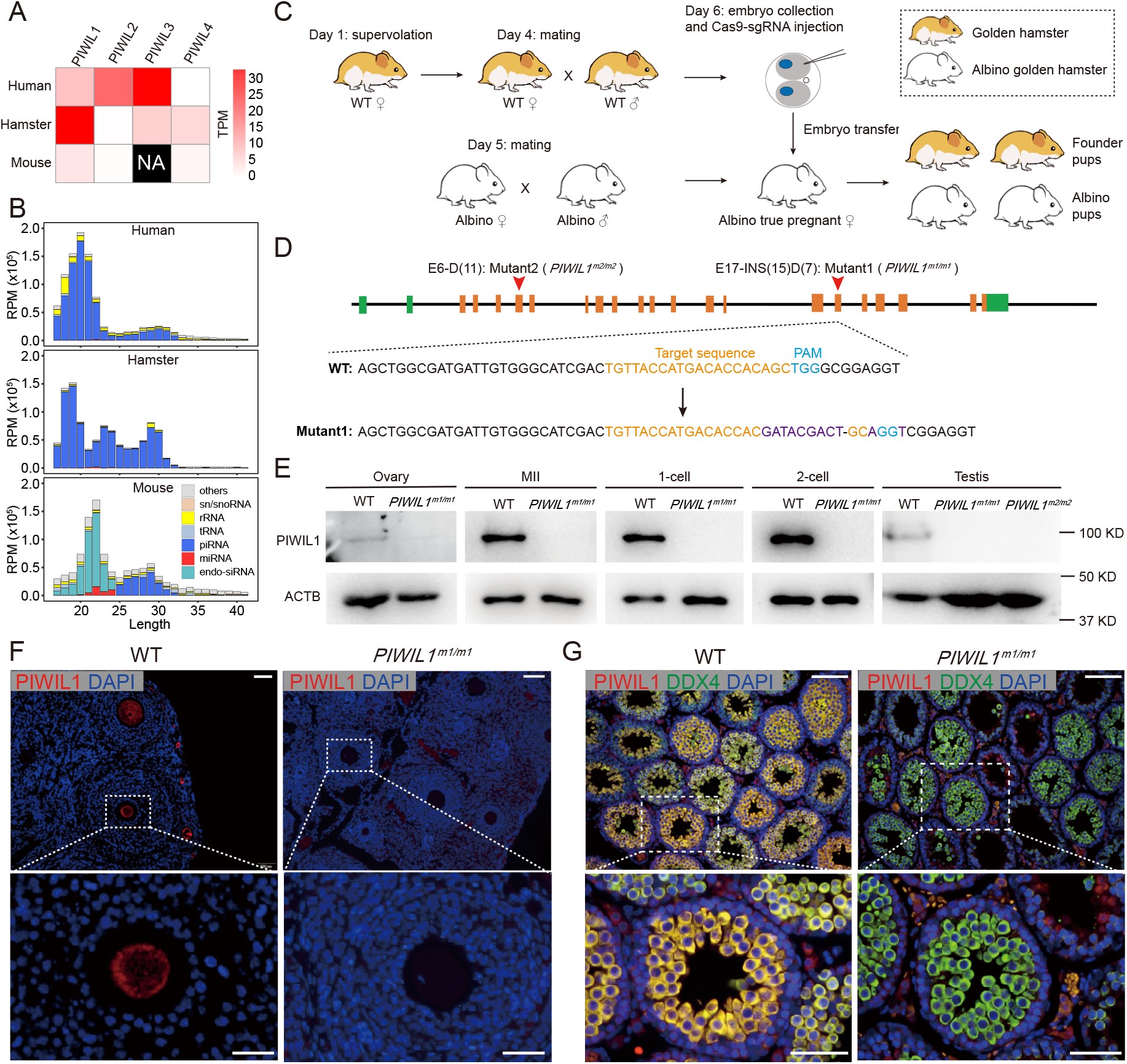
Generation of *PIWIL1* mutant golden hamsters. **(A)** Expression of *PIWIs* in mouse, golden hamster, and human MII oocytes. Expression levels are normalized by transcripts per million (TPM). **(B)** Composition of small RNA categories according to their size distributions in mouse, golden hamster, and human MII oocytes. **(C)** Strategy for the generation of *PIWIL1* mutant golden hamsters by two-cell embryo CRISPR/Cas9 injection. The two-cell embryos from WT golden hamsters were collected and injected with Cas9 mRNAs and sgRNAs, which were then transferred into albino true-pregnant recipient females mated with albino males to obtain *PIWIL1*-edited founder pups. **(D)** Structure of the golden hamster *PIWIL1* gene and generation of *PIWIL1* mutants. Two *PIWIL1* mutant strains were generated. Mutant1 (*PIWIL1*^m1/m1^) contained a 7 nt deletion and a 15 nt insertion in exon 17. Mutant2 (*PIWIL1*^m2/m2^) contained an 11 nt deletion in exon 6. Both of the mutations caused a frameshift in the PIWIL1 mRNA resulting in a premature stop codon. **(E)** Western blots showing the loss of PIWIL1 expression in ovary, MII oocyte, one-cell embryo, two-cell embryo, and testis of *PIWIL1*^m1/m1^ hamsters. ACTB was used as the loading control. **(F-G)** Immunostaining shows loss of PIWIL1 expression in the mutant ovary (F) and testis (G). Scale bars = 100 μm (top); Scale bars = 25 μm (bottom).

## Results

Although CRISPR/Cas9 technology has been used successfully in golden hamsters (*36, 37*), the production of genetically-modified golden hamsters has remained a challenge (*36*). We optimized several critical steps for producing mutant golden hamsters. First, we injected CRISPR/Cas9 into 2-cell embryos, rather than the pronucleus, and then cultured the injected embryos in HEMC9+PVA medium, which significantly increased the survival rate of embryos by more than 50% (*36*, 38, 39) (data not shown). Furthermore, we generated an albino (*Tyr*-deficient) golden hamster strain to simplify the procedure for identifying genome-edited pups by their coat color, while true pregnant females were used as recipients (Fig. s1).

We generated PIWIL1-deficient golden hamster strains by two-cell embryo injection (Fig. 1C) with CRISPR/Cas9 using two sgRNAs targeting exon 6 and exon 17 of *PIWIL1*, respectively. Two *PIWIL1* mutant strains were obtained (Fig. 1D and Fig. s2A) and both mutations caused frame shifts that resulted in premature stop codons in *PIWIL1* mRNAs (Fig. s2B). In homozygous *PIWIL1*-deficient animals, no PIWIL1 protein was detectable in the ovaries, MII oocytes, early embryos, or testes by western blotting (Fig. 1E) and immunostaining (Fig. 1F-G and Fig. s2C-D), thus confirming the complete loss of PIWIL1 expression. Both male and female *PIWIL1* homozygous mutants appeared physiologically normal but were infertile (Fig. 2A). The two independent *PIWIL1* mutants (*PIWIL1*^m1/m1^ and *PIWIL1*^m2/m2^) showed identical results, which ruled out potential off-target effects of Cas9/sgRNA.

**Fig. 2.**
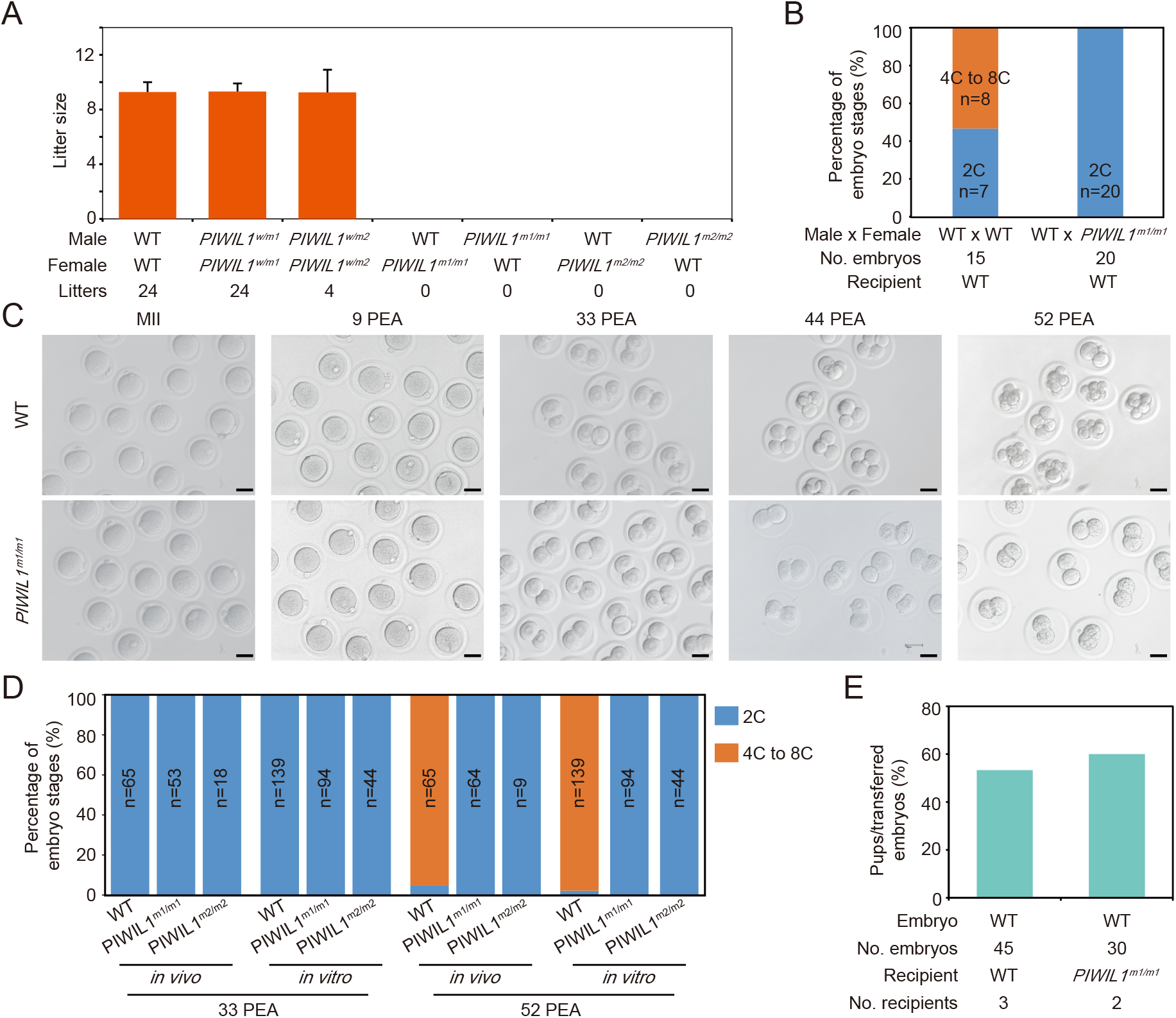
*PIWIL1* deficiency causes infertile in both male and female golden hamsters. **(A)** Fecundity of male and female *PIWIL1* mutant golden hamsters. Error bars indicate standard error of mean (s.e.m). **(B)** Embryogenesis ratio of *PIWIL1*^m1/m1^ embryos transferred to WT recipients. n is the number of embryos at each indicated stage. **(C)** The development of *PIWIL1* mutant embryos was arrested at the two-cell stage *in vivo*. MII oocytes and embryos from the oviducts of WT and *PIWIL1*^m1/m1^ females mated with WT males were collected at 0, 9, 33, 44, and 52 hours post egg activation (PEA). Scale bar = 50 μm. **(D)** Embryonic development of *PIWIL1* mutants was arrested at the two-cell stage *in vivo* and *in vitro*. Embryos from WT or *PIWIL1*^m1/m1^ females mated with WT males were collected at 33 or 52 PEA and their *in vivo* embryogenesis ratio was calculated. For *in vitro* analysis, zygotes from WT and *PIWIL1*^m1/m1^ females mated with WT males were collected at 9 PEA and cultured *in vitro;* embryogenesis ratios were calculated at 33 and 52 PEA. **(E)** The ratio of pups resulting from WT embryos transferred to WT and *PIWIL1*^m1/m1^recipients, respectively.

The testes of PIWIL1 homozygous mutants were substantially smaller than their WT or heterozygous counterparts (Fig. s3A-B). No mature sperm was found in the caput and cauda epididymis of mutants (Fig. s3D-E). Histological examination revealed that round and elongated spermatids were completely absent and that spermatogenesis was uniformly arrested at the pachytene stage (Fig. s3C), which was further confirmed by immunostaining of γH2AX (Fig. s3F) (*24*). These observations indicated that PIWIL1 is essential for spermatogenesis in golden hamsters and that these mutants showed comparable developmental abnormality to *PIWIL1* mutant male mice, as reported previously (*26*).

*PIWIL1* mutant females displayed normal ovarian histology (Fig. s4A), with no obvious differences from the WT in the number of ovulated metaphase II (MII) oocytes (Fig. s4B) or morphology of the MII spindle (Fig. s4C). We found that MII oocytes from *PIWIL1*^m1/m1^ could be successfully fertilized *in vivo* and subsequently form normal pronuclei at 9 hours post egg activation (9 PEA) (Fig. 2C and Fig. s5A). However, in contrast to the WT zygotes, *PIWIL1*^m1/m1^ zygotes were arrested at the two-cell stage *in vivo* (Fig. 2C-D and Fig. s5A). Those arrested two-cell embryos could be maintained up to 52-54 PEA, but beyond that time these embryos appeared to die and degrade in the oviduct. To confirm the embryogenesis defects, zygotes (collected at 9 PEA) were isolated from the oviduct and cultured *in vitro*. A similar two-cell arrest phenotype was observed (Fig. 2D and Fig. s5B) and the embryos broke down after 52 PEA, while WT embryos grew continuously beyond the four-cell stage. Interestingly, the microtubule bridge between two sister cells was absent in *PIWIL1* mutant embryos at 33 to 52 PEA (Fig. s5D), indicating abnormal cell division (*40*). These observations were confirmed in an independent *PIWIL1* mutant stain (*PIWIL1*^m2/m2^) (Fig. s5C), suggesting the embryogenesis deficient phenotype is attributable to PIWIL1.

To rule out the possibility that *PIWIL1* disruption caused defects in the oviduct and ovary that could lead to sterility, we transferred two-cell embryos at 33 PEA from *PIWIL1*^m1/m1^ females mated with WT males into WT recipient females (Fig. s6A). The embryos failed to pass the two-cell stage in these surrogate females (Fig. 2B). Conversely, embryos from WT albino hamsters were transferred into the oviducts of *PIWIL1*^m1/m1^ females (Fig. s6B), and the *PIWIL1*^m1/m1^ surrogate females gave birth to albino hamsters with normal efficiency (Fig. 2E). In addition, we found that *PIWIL1* heterozygotes appeared physiologically normal and gave birth to *PIWIL1*^m1/m1^ offspring at the expected ratio. These results suggested that the loss of maternal PIWIL1 might be the cause of oocyte incompetence and contributed to early embryogenesis arrest.

To further investigate whether a similar phenotype could also be observed upon knockout of other essential genes in the piRNA pathway, we generated mutants of *PLD6 (or ZUCCHINI or MITOPLD)* endonuclease and *MOV10L1* helicase, respectively, which are essential for piRNA biogenesis (*27-30*) (Fig. s7A-B, Fig. s8A-B). The resulting heterozygous *PLD6* and *MOV10L1* mutant males and females showed normal reproductive ability, whereas homozygous *PLD6-* and *MOV10L1-*deficient male and female hamsters were sterile (Fig. s7C-D, Fig. s8C-D). Notably, *PLD6* and *MOV10L1* mutants displayed embryonic arrest at the two-cell stage similarly to the *PIWIL1* mutant (Fig. s7E-F, Fig. s8E-F), thus confirming the indispensable role of the piRNA pathway in female fertility in golden hamster.

Because knockout of *PIWIL1, PLD6*, and *MOV10L1* all resulted in similar phenotypes, we then focused on the *PIWIL1* mutants to investigate the underlying mechanisms that lead to abnormalities in oocytes. The oocytes and embryos at different developmental stages were collected from WT and *PIWIL1*^m1/m1^ females for single-cell small RNA and transcriptome sequencing with exogenous spike-in. The sequencing results revealed the presence of three populations of differently sized piRNAs, including 18-21 nt (19 nt-piRNA), 22-24 nt (23 nt-piRNA), and 28-30 nt (29 nt-piRNA), the relative abundances of which were dynamically regulated during oogenesis and early embryogenesis (Fig. 3A). We characterized these piRNAs and found all of them showed a strong preference for uridine (U) at their 5**’** end and other typical piRNA characteristics (Fig. s9-10)(*7, 8*). Upon NaIO4 treatment, 23 nt-piRNAs and 29 nt-piRNAs remained intact while 19 nt-piRNAs almost completely disappeared (Fig. s10A). These findings indicated that 23- and 29 nt-piRNAs possessed 2**’**-O-methylation at their 3**’** end, while the 19 nt-piRNAs did not, which was consistent with the features of long piRNAs and os-piRNAs previously reported in human and monkey oocytes (*1*). In *PIWIL1* mutants, 23 nt-piRNAs and 29 nt-piRNAs were completely undetectable, while 19 nt-piRNAs and miRNAs were unaffected (Fig. 3A and Fig. s10B). We further sequenced small RNAs associated with PIWIL1 and found that 23- and 29 nt-piRNA both co-purified with PIWIL1 in WT MII oocytes but were absent in PIWIL1-deficient oocytes (Fig. s10C-D), consistent with observations of diminished 23- and 29 nt-piRNAs in the PIWIL1 mutants (Fig. 3A). Furthermore, the 23- and 29 nt-piRNA shared the same piRNA clusters and 5**’** ends (Fig. s11), which supported the likelihood that both 23 nt-piRNAs and 29-nt piRNAs are associated with PIWIL1 in hamster oocytes (*35*). In the testes, only the 29 nt-piRNAs were found to bind with PIWIL1 and these piRNAs were completely absent in *PIWIL1* mutants (Fig. s12), which suggested that PWIL1-associated piRNAs in hamster testes shared similar characteristics and function to their counterparts in mouse testes (*4, 5*).

**Fig. 3.**
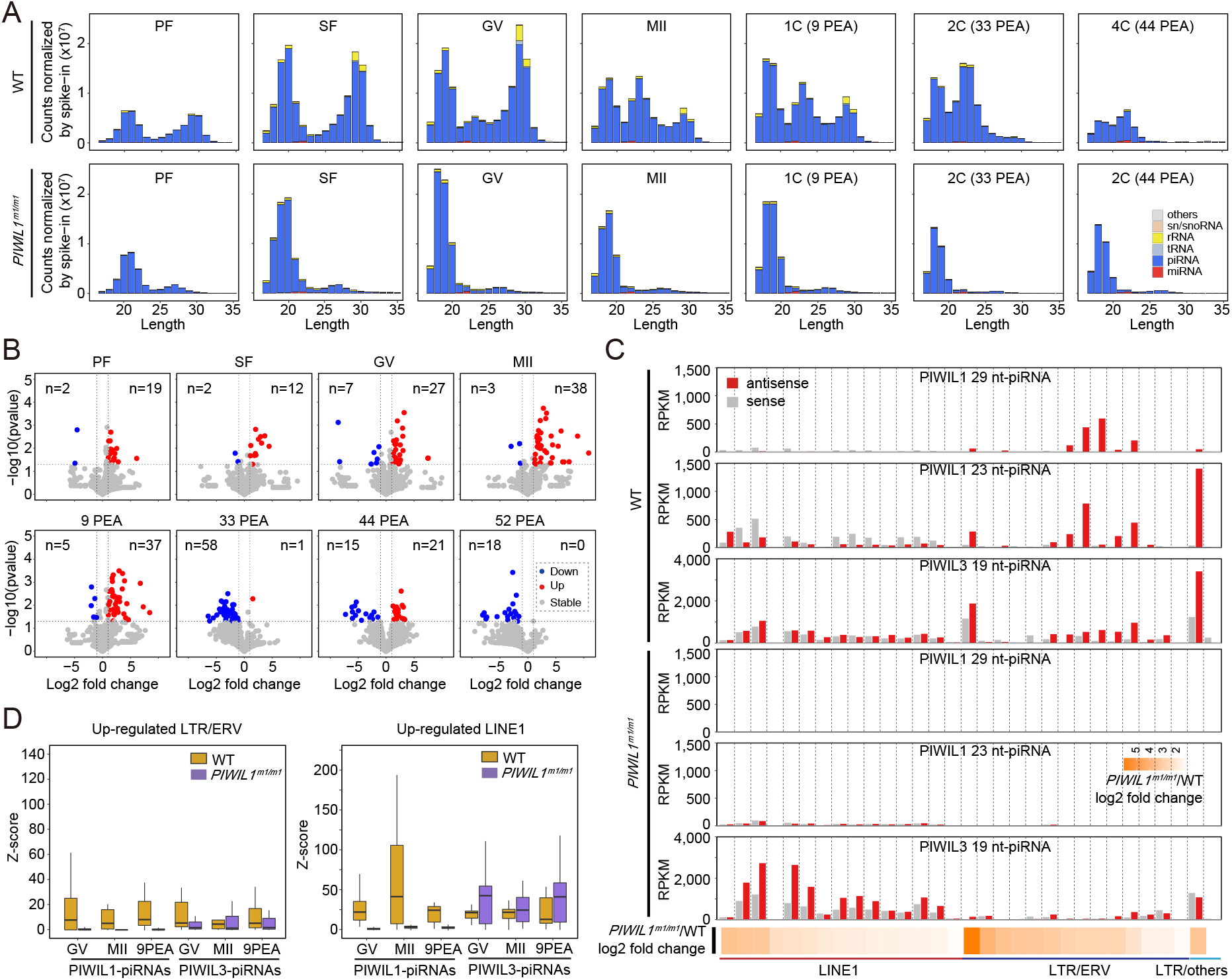
PIWIL1-piRNAs are indispensable for suppression of ERVs but not for LINE1s. **(A)** Composition of small RNA categories according to their length distribution. Early-stage oocytes were collected from the ovaries, and MII oocytes were collected by superovulation. The embryos were collected from the oviducts of WT and *PIWIL1*^m1/m1^ females mated with WT males. The average of 2-6 biological replicates are shown. The small RNA counts were normalized by exogenous spike-in. PF, primary follicle stage oocyte; SF, secondary follicle stage oocyte; GV, germinal vesicle stage oocyte; MII, metaphase II oocyte; 1C, one-cell embryo; 2C, two-cell embryo; 4C, four-cell embryo. **(B)** Differential analysis of consensus TE expression in WT and *PIWIL1*^m1/m1^ oocytes and embryos. The expression of TEs was normalized by exogenous spike-in. The highly significant up- or down-regulated TEs (**≥** 2 folds; p-value < 0.05 determined by Welch two sample *t*-test) are indicated in red or blue, respectively, and the TE number is shown at the top. **(C)** The bar graphs show the expression level (RPKM) of piRNAs mapped to the sense (gray) and antisense (red) direction of each TE family. PIWIL1 29 nt-piRNAs, PIWIL1 23 nt-piRNAs, and PIWIL3 19 nt-piRNAs derived from different TE families are plotted. The significantly up-regulated TE families are listed with fold changes of expression level in *PIWIL1*^m1/m1^ versus WT MII oocytes. The log2 fold change level is indicated as the degree of orange in the heatmap. **(D)** The Ping-Pong signature of PIWIL1-piRNAs (23 nt-piRNA and 29 nt-piRNA) and PIWIL3-piRNAs (19 nt-piRNA) derived from up-regulated LTR/ERV and LINE1 subfamily members in WT and *PIWIL1*^m1/m1^ MII oocytes. The level of Ping-Pong signature is represented by the *Z*-score of 10-nt overlapped piRNAs from opposite strands, with piRNAs having overlaps of different lengths serving as the background. *Z*-score >1.96 corresponds to p-value < 0.05 (*50*).

Approximately 35% of the piRNAs could be mapped to known regions of the genome. Among them, most were mapped to TEs (Fig. s13 and Fig. s14), which was consistent with the conserved function of piRNAs in silencing TEs (*7, 8*). Our transcriptome data from oocytes and embryos at different developmental stages revealed that TEs accumulated during follicle development and started to decrease after the germinal vesicle (GV) stage in WT (Fig. s15A), possibly due to suppression by the piRNA pathway. Intriguingly, a minor wave of TE activation was observed in one-cell and two-cell embryos, reminiscent of similar observations reported during early embryogenesis of other mammals (*41, 42*). In the *PIWIL1* mutant, TE expression was significantly up-regulated, especially from the GV oocyte to one-cell embryo stages (Fig. 3B). This trend was sustained until 33 PEA, at which point *PIWIL1*-mutant embryos underwent developmental arrest and subsequent degradation. In particular, ERVs were the most highly up-regulated TEs, followed by LINE1s (Fig.s15B). The ERV-related PIWIL1-piRNAs showed an obvious bias to the antisense of ERV mRNAs (Fig.3C and Fig. s16) and a strong Ping-Pong signature (Fig. 3D and Fig. s17), which suggested that involvement of PIWILI cleavage guided by antisense piRNAs in silencing ERVs. Notably, the ERV-related PIWIL3-piRNAs were almost completely depleted in oocytes and embryos upon PIWIL1 disruption (Fig. 3C, Fig. s16B and Fig. s17), implying that generation of ERV-related PIWIL3-piRNAs, to some extent, depends on PIWIL1, possibly through inter-Ping-Pong crosstalk between two different PIWIL proteins (*43, 44*). These results highlight the non-redundant and essential role of PIWIL1 in silencing ERVs.

For LINE1s, both PIWIL1- and PIWIL3-piRNAs showed an obvious Ping-Pong signature (Fig. 3D). Intriguingly, the abundance of LINE1-related PIWIL3-piRNAs, as well the Ping-Pong signature, were increased following *PIWIL1* knockout (Fig. 3C-D, Fig. s16), which was potentially caused by high accumulation of TE mRNAs and/or primary piRNA transcripts in the *PIWIL1* mutants. These results suggested that both PIWIL1 and PIWIL3 were involved in LINE1 silencing and exhibited redundant functions. The differential activity in silencing ERVs and LINE1s by PIWIL1 and PIWIL3 might be attributable to the preferential interaction of individual PIWI proteins with the pre-piRNA processing complexes, which warrants further investigations.

In light of these findings, we speculated that the accumulation of TEs might not be the exclusive reason for the arrest of two-cell stage embryos in *PIWIL1* mutants. Principal component analysis (PCA) for gene expression between stages showed that the oocytes and 1-cell embryos of mutant and WT grouped tightly and were distinguishable from other stages, whereas *PIWIL1*-mutant embryos after 33 PEA clustered separately from WT (Fig. 4A and Fig. s18). Developmental trajectory assays showed that *PIWIL1*-mutant embryos branched off toward a different cell fate from that of WT embryos after 33 PEA (Fig. 4B), which became developmentally incompetent and died within 1-2 days *in vivo* and *in vitro*. Quantification of mRNAs showed 254 and 1186 differentially expressed genes (DEGs) in 1-cell and 2-cell embryos of *PIWIL1*-mutants, respectively. Notably, the DEGs significantly elevated in WT embryos were rarely up-regulated in *PIWIL1* mutant embryos (Fig. 4C and Fig. s19), indicating that minor and major zygote gene activation (ZGA) was severely inhibited in *PIWIL1*-mutants.

**Fig. 4.**
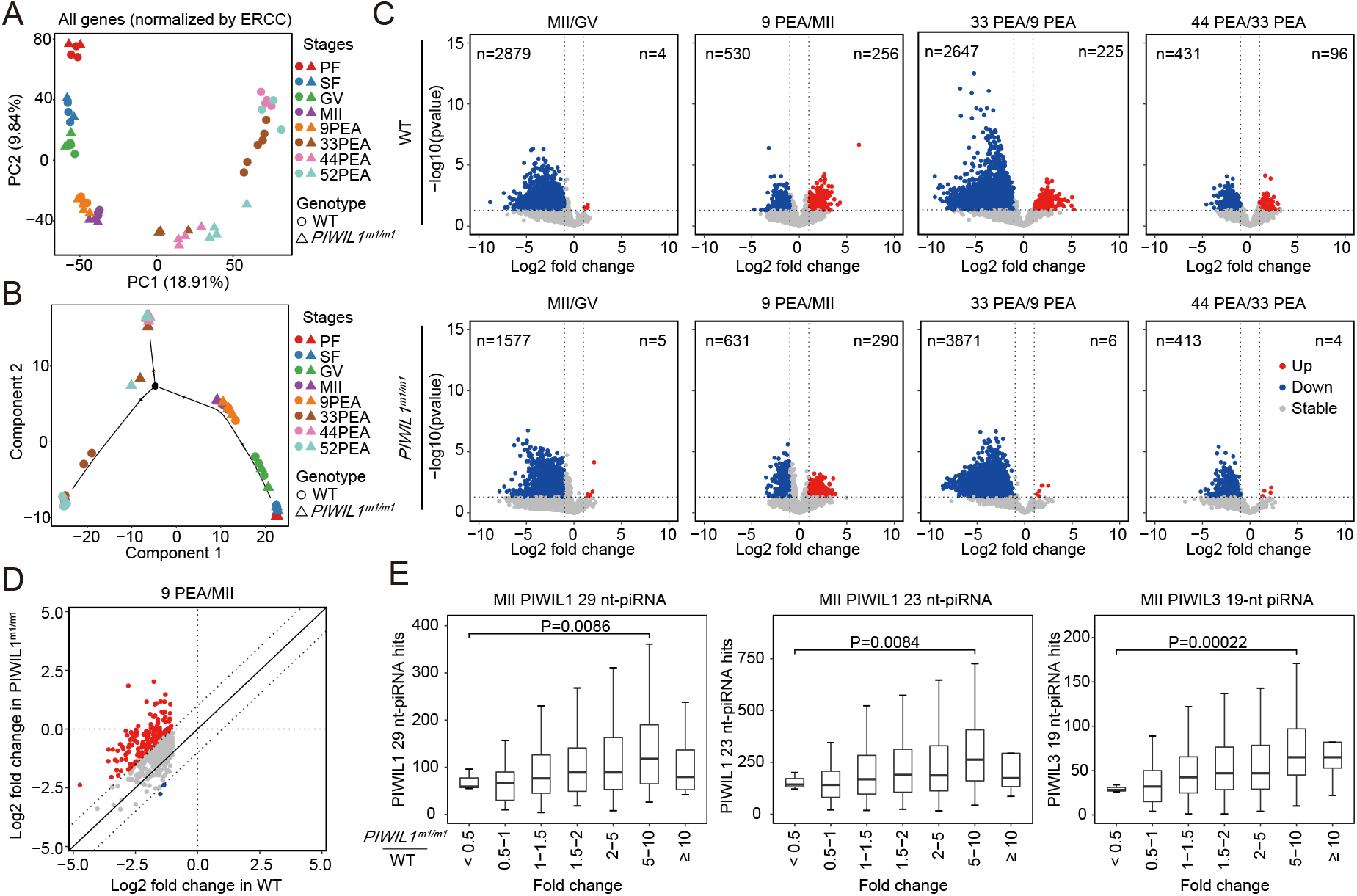
PIWIL1-piRNAs participate in the degradation of maternal mRNAs. **(A)** PCA analysis of all genes expressed in the oocytes and embryos at different stages. The circles represent WT, and triangles represent *PIWIL1* mutants. The gene expression levels were normalized by exogenous spike-in. **(B)** Pseudotime analysis of all gene expression by Monocle reveals a bifurcation between the developmental trajectories of WT and *PIWIL1*^m1/m1^ embryos into two distinct branches after 9 PEA. **(C)** Volcano plots show the differentially expressed genes in oocytes or embryos between GV and MII stages, MII stage and 9 PEA, 9 PEA and 33 PEA, 33 PEA and 44 PEA. The highly significant up- or down-regulated genes (**≥** 2 folds; Welch two sample *t*-test, p-value < 0.01) are indicated in red or blue, respectively, with gene number shown at the top. **(D)** Degradation of some maternal mRNAs is inhibited between the MII stage to 9 PEA in *PIWIL1* mutants. X- and Y-axes represent the log2 fold change of gene expression in WT (X-axis) and *PIWIL1*^m1/m1^ (Y-axis) at 9 PEA versus MII. **(E)** The number of targets on the mRNAs matched to significantly down-regulated piRNAs after PIWIL1 knockout was correlated with the level of mRNA up-regulation. The level of target mRNA up-regulation was calculated as the ratio of fold changes in embryos at 9 PEA to MII oocytes between *PIWIL1*^m1/m1^ and WT samples.

Initiation of ZGA is known to coincide with the degradation of maternal mRNAs, which is started in GV oocytes to remove repressive factors and to enable zygotic transcription, as well as to permit the establishment of embryonic patterning (*45*), raising the possibility that the inhibition of ZGA in *PIWIL1-*deficient embryos might be trigged by impaired decay of maternally deposited transcripts, with deleterious consequences. To examine this hypothesis, we investigated the relative mRNA abundance in each stage and found that the decay rate of many rapidly degraded mRNAs dramatically decreased in *PIWIL1* mutants during the MII oocyte to 1-cell transition (Fig. 4D).

Previous studies showed that piRNAs affect the expression of mRNAs by two distinct mechanisms: one relies on PIWIL1 slicer activity to cleave almost perfectly complementary targets; the other mechanism is to trigger deadenylation of partially complementary targets via recruitment of the deadenylation complex (*9, 10, 19, 46*). We observed a significant enrichment of target sequences that were partially complementary to PIWIL1-piRNAs on those mRNAs with impaired degradation. The number of targets correlated with the changes in the degradation rate of mRNAs upon *PIWIL1* deletion (Fig. 4E), as did the abundance of piRNAs (Fig. s20A), which suggested the involvement of PIWIL1 and its associated piRNAs in controlling maternal mRNA stability. Notably, the PIWIL1 deficiency in MII oocytes also resulted in impaired degradation of many predicted target mRNAs of PIWIL3-piRNAs whose abundance was affected in the absence of PIWIL1 (Fig. s14, Fig. s20B-C). These results implied that PIWIL1 might also indirectly regulate additional maternal mRNA degradation through PIWIL3 and its associated piRNAs.

## Discussion

Mice are the most commonly used animal model for studying gene function and human disease. However, growing evidence shows that mice may not be as widely representative of mammals in some genetic and physiological aspects as previously thought (*34*). piRNAs appear to be one such case. Previous studies have shown that PIWIs and piRNAs are essential for female fertility in flies and fish (*20-23*), but dispensable for female fertility in mice (*24-30*), a discrepancy likely due to the lack of *PIWIL3* in the mouse genome, as well as mouse-specific expression of endo-siRNAs that functionally overlap with piRNAs for silencing TEs in oocytes (*31, 47*). In this study, we showed that the piRNA pathway is essential for female fertility in golden hamsters. In addition to their highly conserved function in silencing TEs, we show that piRNAs also function in the regulation of maternal mRNA stability in oocytes and embryos, and that disruption of the piRNA pathway leads to gene expression dysregulation in oocytes and failure to initiate ZGA in embryogenesis. Previous studies have shown that in *Drosophila, Aubergine* (*PIWI* homolog) and its associated piRNAs recruit Smaug and the CCR4 deadenylation complexes to eliminate specific maternal mRNAs in the early embryo (*19*); and in *Aedes aegypti*, satellite repeat-derived piRNAs degrade maternally deposited transcripts in the zygote (*17*), indicating that the role of piRNAs in maternal mRNA clearance is conserved across vertebrate and invertebrate species. During the preparation of this report, two other groups have described similar observations (albeit in non-peer-reviewed preprints) corroborating the indispensable functions of *PIWIL1* and *MOV10L1* in female fertility in golden hamster (*48, 49*). Given the small size, ease of maintenance, high reproductive performance, and most importantly, similar patterns of PIWIs and piRNAs expression in oocytes compared to humans, we propose an expanded role for golden hamster as an instructive and tractable representative model for mammalian reproduction. In particular, the investigation of piRNA-related functions in females can yield valuable insights with potential clinical implications for fertility-related disorders.

## Acknowledgment

We thank all members of Ligang Wu**’**s and Jianmin Li**’**s laboratory for their discussion and comments on this project. We thank Zhixue Li for bioinformatics analyses, Yuanyuan Song for library construction. We thank Yaochen Xu for his assistance with high-performance computing. We are grateful to the HPC storage and network service platform of SIBCB for supplying the computing resources. We thank Dr. Haruhiko Siomi (Keio University School of Medicine, Tokyo, Japan) for sharing unpublished data. We are also grateful to Ying Huang, Mofang Liu, and Yang Yu for their critical comments on the manuscript.

