## Supplementary materials for "piRNA pathway is essential for generating functional oocytes in golden hamster"

**List of supplementary materials**

Materials and Methods

Supplementary Figures s1-s20

Supplementary data Table s1

### Materials and Methods

#### Animals and ethics statement

Golden (Syrian) hamsters were purchased from Vital River (Charles River, China) and Liaoning Changshen Biotechnology Co. Ltd, and maintained under 14-h light/10-h dark cycles (dark from 20:00-6:00). All experiments were approved by the Nanjing Medical University Institutional Animal Care and Research Committee (IACUC code: 1709024).

#### Collection of oocytes and embryos

Golden hamster follicles and germinal vesicle (GV) stage oocytes were collected from ovaries as described previously (*1, 2*). Female golden hamsters of 6-8 weeks old were induced to superovulation by injection of 10-30 IU pregnant mare serum gonadotropin (PMSG, Ningbo Second Hormone Company, Cat# 110254564) according to their body weight on the first day of their estrus cycle. For MII oocyte collection, hamsters were injected with 15 IU human chorionic gonadotropin (hCG, Ningbo Second Hormone Company, Cat# 110251281) at 56-58 h after PMSG injection and the MII oocytes were collected from oviducts at 16-18 h after hCG injection. To obtain embryos *in vivo*, superovulated female hamsters were mated with males on the fourth day after PMSG injection, and embryos were collected from oviducts at 9 (one-cell embryo), 33 (two-cell embryo), 44 (two- or four-cell embryo), and 52-54 (four-cell embryo) h post egg activation (PEA). To obtain embryos *in vitro*, zygotes were collected from oviducts of mated females at 9 PEA and cultured in mHEMC9+PVA medium under a humidified 10% $CO_2$ environment at 37.5°C and then embryos were collected at 33, 44, and 52-54 PEA. All oocytes and embryos were manipulated and observed under red light in a dark room, and the microscope was equipped with a red-light filter.

#### Generation of mutant golden hamsters

sgRNAs were designed using CRISPRdirect (http://crispr.dbcls.jp/) and CRISPOR (http://crispor.tefor.net/). The complementary target oligonucleotides were cloned into a pUC57-T7-sgRNA-trcRNA vector through *Bsa*I restriction site. The sequences containing T7 promoter and sgRNA were PCR-amplified and used as templates to produce sgRNAs by *in vitro* transcription using HiScribe T7 High Yield RNA Synthesis Kit (NEB, Cat# E2040S). The full-length open reading frame (ORF) of Cas9 with T7 promoter was PCR amplified from a pX330-U6-Chimeric_BB-CBh-hSpCas9 plasmid (from Feng Zhang (Addgene plasmid #42230; http://n2t.net/addgene:42230; RRID: Addgene_42230)) (*3*) and used as templates for *in vitro* transcription with mMESSAGE T7 ULTRA Transcription Kit (Invitrogen, Cat# AMB1345). All sgRNAs and Cas9 mRNAs were purified by lithium chloride precipitation and stored at -80°C.

Microinjection was performed as described previously (*4*) with several modifications. We injected CRISPR/cas9 mRNA and sgRNA into two-cell embryos (two-cell embryo CRISPR/Cas9 injection method, 2CEC injection) instead of pronuclei. We also changed the culture medium from HEMC9+HAS to mHEMC9+PVA to increase the survival rate of embryos; and changed the culture conditions from a humidified 10% $CO_2$, 5% $O_2$ environment to a humidified 10% $CO_2$ environment, which had no effect on the survival rate of embryos but simplified the culture conditions. To generate genome modified golden hamsters, two-cell embryos were collected as described above and injected with Cas9 mRNA, sgRNA, and/or ssDNA donor in a red-light room under a microscope with a red filter. Injected embryos were cultured in mHECM9+PVA medium under a humidified 10% $CO_2$ environment at 37.5°C for 15-30 min, and transferred to the oviducts of 0.5 d pregnant or pseudo-pregnant recipients, with 15-30 embryos for each recipient. Founder pups were mated with wild-type males and females to produce the F1 generation. Genotyping of F1 pups was performed using a one-step mouse genotyping kit following the manufacturer's instructions (Vazyme, Cat# PD101-01). All the oligo sequences are listed in Supplementary Table S1.

**Western blot analysis**

Fifty to one hundred oocytes and embryos (one-cell or two-cell) were collected as described above for western blotting. Testis and ovary tissues were acquired from 8-week-old golden hamsters. Total proteins were extracted in 95% Laemmle sample buffer, separated on 10% SDS-PAGE using a

Mini-PROTEAN Tetra Cell System (Bio-Rad), and electrophoretically transferred to PVDF membranes. Anti-β-actin (ACTB) mouse monoclonal antibody (Proteintech, Cat# 66009-1-lg, 1:2,000) and anti-PIWIL1 rabbit polyclonal antibody (1:1,000) (*5*) were used as primary antibodies and incubated with the membranes at 4°C for 12 h. After washing twice with PBST, the membranes were incubated with HRP-linked goat anti-rabbit (Thermo Fisher, Cat# 31466, 1:5,000) or HRP-linked anti-mouse IgG (Thermo Fisher, Cat# 31430, 1:5,000) at room temperature (RT) for 1 h before visualization using an ECL Detection kit (GE Healthcare, USA).

**Histological analysis**

Testis and ovaries were fixed in Bouin's fixative (Sigma, Cat#MBD1105) overnight at 4°C, embedded in paraffin, and sectioned at 5 μm. For periodic acid-Schiff (PAS) staining, sections were deparaffinized and rehydrated and then stained with PAS. Slides were mounted in neutral resins and images were acquired on a BX53 wide-field microscope (Olympus).

**Immunofluorescence analysis**

Testis and ovaries were fixed in 4% paraformaldehyde (PFA) overnight at 4°C followed by embedding in paraffin and sectioning at 5 μm. Sections were deparaffinized and rehydrated with citrate buffer (pH 6.0) for antigen retrieval. Following blocking with 2% bovine serum albumin (BSA) and 4% normal donkey serum in phosphate-buffered saline (PBS) overnight at 4°C, the sections were treated with anti-PIWIL1 polyclonal antibody (1:200) (*5*), anti-DDX4 monoclonal antibody (1:200) (Abcam, Cat# ab27591), or anti-γH2AX (1:200) (Millipore, Cat#05-636) for 2 h at 25°C. The slides were washed twice with PBS containing 0.2% Tween-20 (PBST), and then treated with Alexa488-conjugated donkey anti-mouse IgG (1:1,000) (Molecular Probes, Cat#A21206) or cy3-conjugated donkey anti-rabbit IgG (1:1,000) (Jackson ImmunoResearch, Cat#711-165-152) as the secondary antibody for 1 h at 25°C. After washing twice with PBST, the sections were stained with 1 μg/ml DAPI (SIGMA, Cat# HT10132). Fluorescent images were captured using a BX53 fluorescence microscope (Olympus).

Oocytes and embryos were collected *in vitro* or *in vivo* and fixed with 4% PFA, permeabilized with 0.2% Triton X-100 in PBS, and then blocked with PBS containing 1% BSA and 2% normal donkey serum overnight at 4°C. The oocytes and embryos were incubated with anti-PIWIL1 antibody (1:200) (*5*), anti-tubulin FITC antibody (Sigma, Cat# F2168, 1:500), or rhodamine-phalloidin (US Everbright, Cat# YP6003, 1:100) for 2 h at RT. For PIWIL1 detection, samples were further incubated with a secondary antibody of Alexa594-conjugated donkey anti-rabbit IgG (Invitrogen, Cat# A21207, 1:200) for 1 h at RT. Nuclei were stained with DAPI. Laser confocal scanning images were captured with an LSM 710 confocal laser scanning microscope (Zeiss).

**Single-cell small-RNA library construction**

Single oocyte or embryo small RNA libraries were constructed as previously described (*5*) with several modifications. Briefly, the single oocytes or embryos were incubated at 72°C for 3 min to release and unfold the small RNAs. After 3' adapter ligation, 5U of lambda exonuclease and 25U of 5' deadenylases, which performed better than the combination of RecJf and 5' deadenylases, were used to remove the excess 3' adaptor. Then, a 5' adapter was ligated and small RNAs were reverse transcribed. Pre-amplification was performed and then 1 μl of amplified product was used as a template for final amplification. The amplified libraries were separated on a 6% polyacrylamide gel and 130-160 bp DNA fragments were sliced to recover the library of small RNAs. All libraries were sequenced at 2x150 bp using a HiSeq X Ten instrument (Illumina, USA).

**Small-RNA spike-in information**

Three un-methylated and three methylated small-RNA spike-ins were synthesized (Integrated DNA Technologies, IDT) and pooled. Spike-ins at $1x10^{-10}$ pmol were added into the reaction for single oocytes or embryos. The sequence of each spike-in can be found in a previous study (*5*).

**Single-cell mRNA library construction**

The procedure was performed following the previously described Smarter-seq2 method with some modifications (*6*). In brief, single oocytes or embryos were incubated at 72°C for 3 min to release the RNAs. For every single oocyte or embryo, 0.125 μL 1/100,000 dilution of the ERCC RNA Spike-In Mix (Invitrogen, Cat# 4456740) was used. Following reverse transcription and PCR pre-amplification, cDNA libraries were purified by Agencourt Ampure XP beads (0.8:1 ratio) (Beckman

Coulter, Cat#A63881) and 2 ng of purified cDNA was used for a tagmentation reaction with Tn5 transposase as described previously (*7*). Amplified libraries were purified by Agencourt Ampure XP beads (1:1 ratio) and sequenced at 2x150 bp on a HiSeq X Ten instrument (Illumina, USA).

**Immunoprecipitation**

Ten MII oocytes of WT and *PIWIL1* mutant golden hamster were lysed with lysis buffer composed of 50 mM Tris-HCl (pH 7.4), 150 mM NaCl, 1 mM EDTA, 0.5% NP-40, 0.5 mM DTT, 0.1 U/μl RNase Inhibitor (Thermo Fisher Scientific, Massachusetts, USA) and 0.4 U/μl Proteinase inhibitor cocktail (Sigma, USA) on ice for 10 min. Dynabeads Protein G (Thermo Fisher Scientific, Massachusetts, USA) were washed once with lysis buffer and coupled with 1.5 μg of homemade anti-PIWIL1 or rabbit IgG (Millipore) at RT for 40 min. Oocyte lysates were mixed with antibody-coupled beads and rotated gently at 4°C for 5 h. The beads were then washed three times with lysis buffer and once with PBS. Total RNA from the beads was eluted by adding 4 μl of water and incubating at 72°C for 5 min. The supernatant containing RNA was recovered on a magnetic holder and then used for cDNA library construction. For testis, fresh tissues were homogenized in lysis buffer using Precellys 24 tissue homogenizer (Bertin, USA) and immunoprecipitation was performed as described above. Small RNA libraries were constructed using the single-cell small RNA library construction method as described above.

**$NaIO_4$ oxidation of oocyte and testis RNAs**

Total RNAs from ten MII oocytes of WT and *PIWIL1* mutant golden hamster were extracted with TRIzol Reagent (Ambion, USA) after adding $1 \times 10^{-8}$ pmole of spike-in oligos. The pellets were dissolved in 4 μl of water. Half of the RNA was saved as a non-oxidation control and the remaining RNA was treated with an oxidation reaction mixture containing 1.5 μl of 100 mM $NaIO_4$ (Sigma, USA), 2 μl of 5× borate buffer (150 mM borax, 150 mM boric acid, pH 8.6) and 4.5 μl of water. The reaction mixtures were incubated in the dark for 30 min at RT and RNAs were precipitated by adding 600 μl of 100% ethanol, 25 μg of linear acrylamide, and 30 μl of sodium acetate (3 M, pH 5.2), then stored at -30°C for 30 min before centrifugation. After washing twice with 75% ethanol, pellets were dissolved in 2 μl of water and ready for cDNA library construction. For the testis, 40 ng of total RNAs were mixed with 1 × $10^{-8}$ pmole of spike-in oligos in a total volume of 4 µl. Half of the RNA mixture was saved as a non-oxidation control, and the remaining RNA was treated with an oxidation reaction mixture and precipitated as described above. All small RNA libraries were constructed using the single-cell small RNA library construction method as described above.

**Categorization of small RNAs**

The small RNA libraries were sequenced on a HiSeq X Ten instrument (Illumina, USA) with 2 × 150 run cycles. Raw read1 fastq was used to identify small RNAs, which were pre-processed using the fastx_toolkit. After quality filtering, sequencing reads were clipped from the 3' adaptor allowing a minimum match of 10 nt from the 5' end. Reads unable to match the adaptor sequence or with lengths shorter than 17 bp were discarded, and the remaining reads were mapped to the golden hamster genome (MesAur1.0) by bowtie with parameters –k=100 –v=0 (*8*). If the sequences mapped to the genome more than 100 loci, we randomly output 100 of all mapped loci. The mapped genome sequences were further aligned to miRNAs, tRNAs, rRNAs, snoRNAs, and snRNAs by bowtie with parameters –a –v=0 --norc.

The remaining 17–32 nt sequences were used to identify piRNAs following a previously described method with slight modifications (*9*). The clustering parameters were determined as MinReads = 4 and Eps = 2500 bp by running a series of k-dist analyses of our data with different Eps and MinReads. All of the clusters that satisfied these parameters were considered as piRNA cluster candidates and any overlapping clusters were merged into one cluster. The 17-32 nt unknown reads located in these clusters were defined as piRNA candidates. The small RNAs were mapped in the following order: miRNA, tRNA, rRNA, snoRNA, snRNA, and piRNA. The reads in each library were normalized with sequencing depth or the count of exogenous spike-ins.

***De novo* miRNA identification and expression profiling**

We constructed and sequenced the small RNA libraries of liver, lung, intestine, testis, and ovary from adult male golden hamsters. The reads mapped to the genome of the golden hamster were combined to predict miRNAs *de novo* by miRDeep2 with default parameters. Only the read counts of star sequences more than 3, and mature sequences more than 10 were classified as miRNA precursors (pre-miRNAs), and others were dropped. Only reads that exactly matched the 5' start site of identified miRNAs and 3' ends with ≤2 nt deletions or additional sequences derived from pre-miRNAs were counted for miRNA expression level.

**Data analysis of single-cell RNA-seq and differential expression analysis**

The cDNA libraries were sequenced on a HiSeq X Ten instrument (Illumina, USA) with 2 × 150 run cycles. The raw paired-end fastq reads were clipped off the adaptors and quality filtered by trim_galore with parameters --quality 20 --stringency=1 –length=35 --paired (http://tophat.cbcb.umd.edu/). The remaining fastq reads were mapped to the genome or identified transposable element (TE) representative sequences by STAR with parameters --winAnchorMultimapNmax 10000 --outFilterMultimapNmax 10000. The reads uniquely and multiply mapped with genomic copies less than 10,000 were used for subsequent analysis. The expression level of each gene and retrotransposon was normalized by ERCC spike-ins. Differentially expressed genes or retrotransposons were identified by applying combined thresholds based on fold change (≥ 2 or ≤ 0.5), and Welch two sample *t*-test (p-value < 0.01 for gene, p-value <0.05 for retrotransposon).

**Principal component analysis (PCA) and pseudo-time analysis**

PCA was performed based on all of the genes expressed in each sample by Seurat v3 with default parameters (*10*). Pseudo-time analysis was performed based on all of the genes expressed in each sample by Monocle v2 (*11*) with default parameters.

***De novo* identification of transposable elements (TEs) in the golden hamster**

RepeatModeler2 (*12*) with default parameters was used to predict and classify TEs. The resulting repeat models were searched against the GenBank non-redundant protein database (e-value <10−5) using Blastx (v2.2.28) (*13*) to exclude potential protein-coding genes. Annotations of all predicted representative TEs were established in the golden hamster genome (MesAur1.0) by RepeatMasker (version 4.1.1) (*14*) with the parameters -e crossmatch −no_is −gff and −lib.

**piRNA genomic annotation**

For different subgroups of TEs or genes, piRNA reads in the sense and antisense direction were counted in order as ERV, other_LTRs, LINE/L1, retrotransposon/L1-dep, LINE/L2, SINE/Alu, SINE/B2, SINE/others, Low complexity/Simple_repeat/Satellite (LSS), DNA transposon, other_repeats, protein-coding exon, pseudo-gene exon, other_exonic, intron and unknown.

**Expression profiling for consensus TE-derived piRNAs**

Identified piRNA sequences were mapped to the consensus TE sequences using bowtie allowing up to 2-nt mismatches in two directions (*8*), and the RPKM (reads per kilobase per million mapped reads) or RPM (reads per million mapped reads) of each TE-derived piRNA was calculated. If a piRNA could be mapped to different members of a TE subfamily, this piRNA was counted for each TE member.

**piRNA target prediction and regulatory function analysis**

piRNA target prediction was performed by requiring perfect pairing between a target and nucleotides 2–8 from the piRNA 5'end. If a gene had several transcript variants, the target number of the longest variant was counted. For the analyses shown in Figure 4-E and related Supplemental Figures, genes were classified into seven groups based on the extent of up-regulation of expression level during MII oocyte development into the 9 PEA stage in *PIWIL1-KO* versus wild-types (<0.5, 0.5-1, 1-1.5, 1.5-2, 2-5, 5-10 and ≥10), while for 1.5-2, 2-5, 5-10 and ≥10 groups, only those genes with significant change (Welch two sample *t*-test p<0.01) in expression level were used for analysis.

**Sources of annotations**

Golden hamster genome annotation (MesAur1.0) was downloaded from Ensembl release 102 (https://asia.ensembl.org/index.html). Gene annotations were also downloaded from Ensembl. Functional RNA from golden hamster was retrieved from the following databases: tRNAs, RNACentral v17 (https://rnacentral.org/); 5S and 5.8S from Ensembl; snoRNAs and snRNA from Ensembl. Moreover, to fully classify the functional RNA-derived small RNAs, we combined the known functional RNA sequences annotated in mouse and rat genomes with related annotations in the golden hamster. For mice and rats, 18S, 28S, and 45S were downloaded from NCBI nucleotide database (https://www.ncbi.nlm.nih.gov/nucleotide/); tRNA was retrieved from gtRNAdb (http://gtrnadb.ucsc.edu/) (*15*); others were retrieved the same way as in golden hamster.

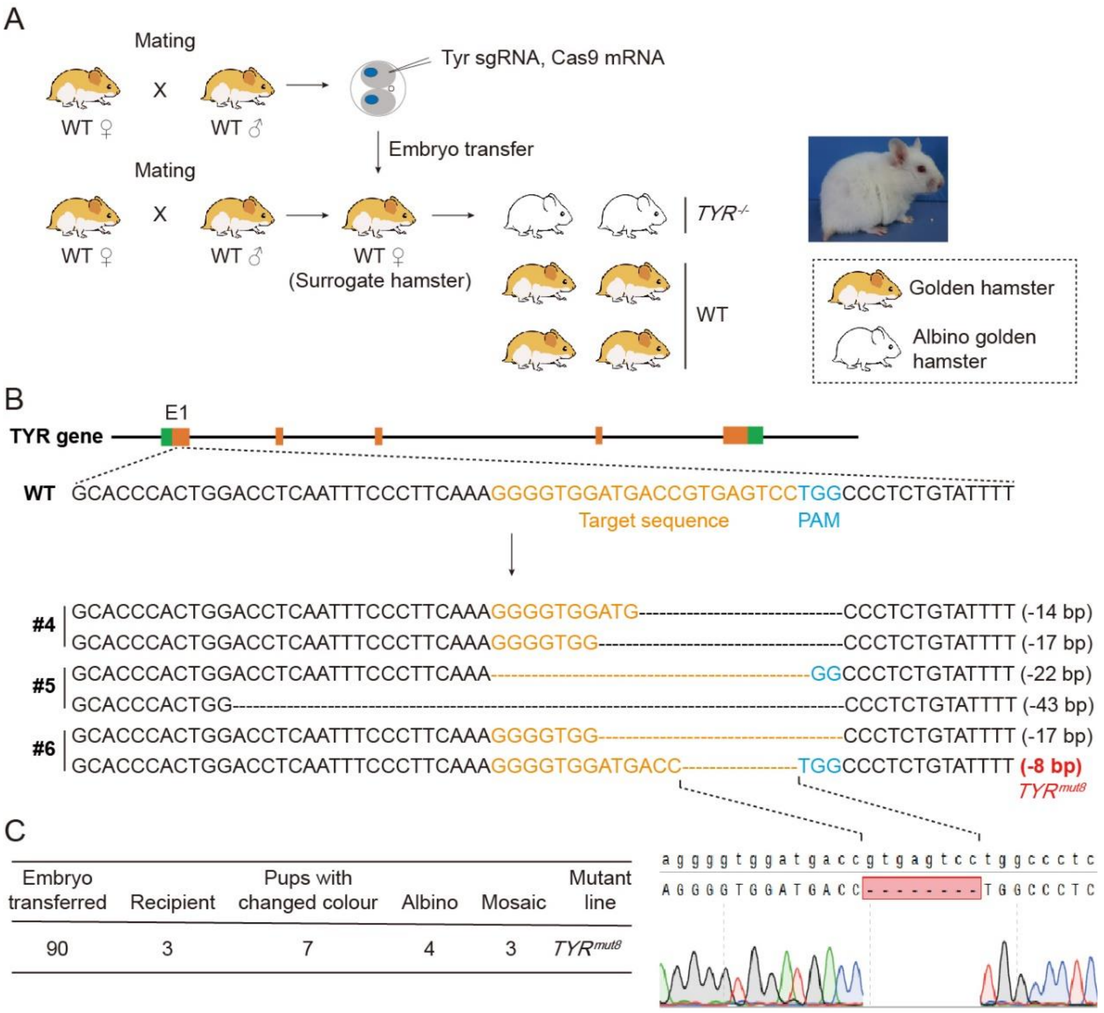


**Fig. s1 Generation of albino golden hamsters**

**(A)** Strategy for the generation of albino (*TYR* mutants) golden hamsters. Two-cell embryos were injected with CRISPR/Cas9 and sgRNAs and transferred to naturally pregnant recipients. **(B)** Diagram of the golden hamster *TYR* gene and the resulting *TYR* mutants. The sgRNA was designed to target Exon 1. **(C)** Production of albino golden hamster lines. Albino golden hamster lines were established by 7 mutant founders.

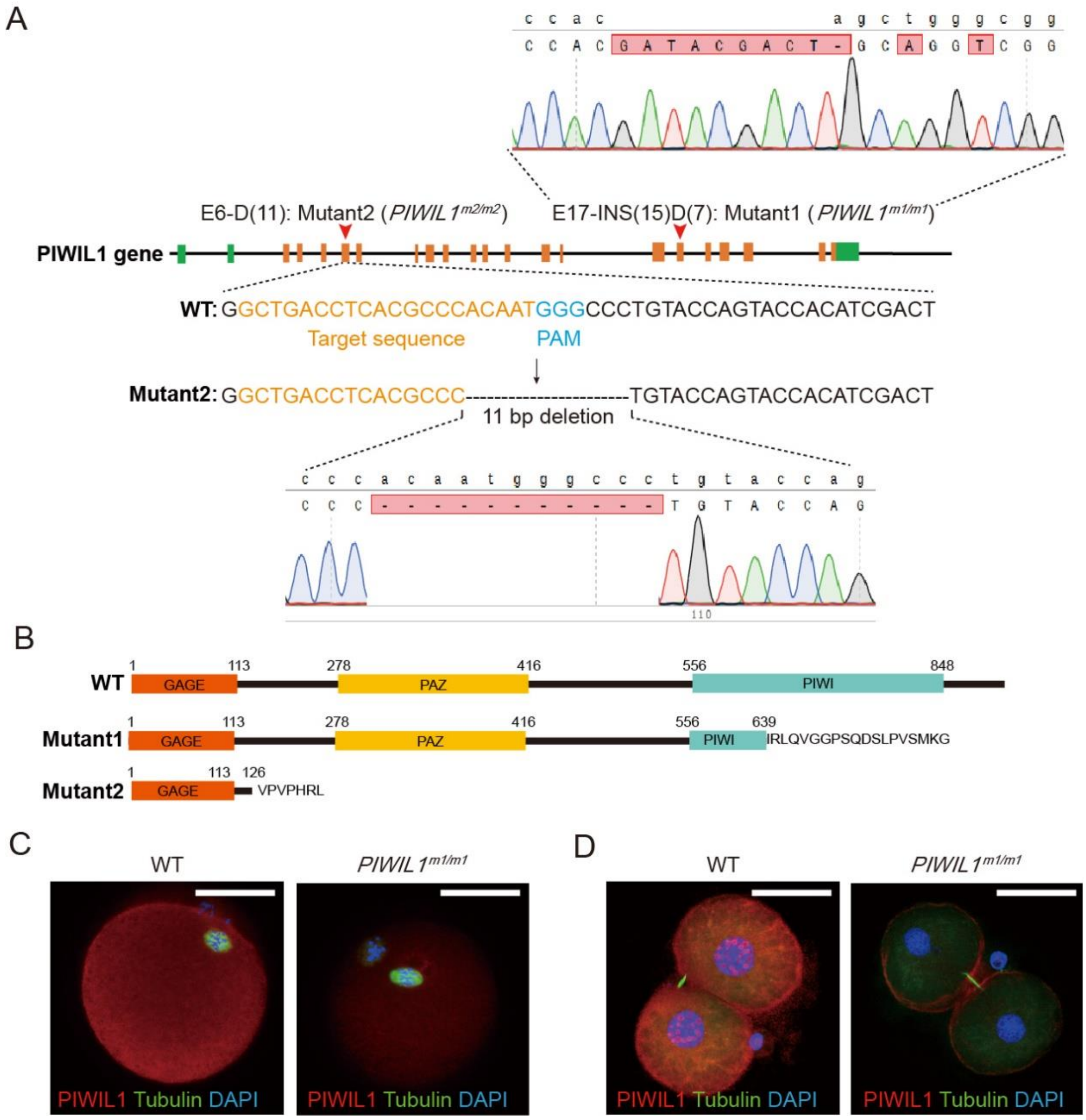


**Fig. s2 Generation of *PIWIL1* mutant golden hamsters**

**(A)** Strategy for the generation of *PIWIL1* mutant golden hamsters by CRISPR/Cas9. **(B)** Diagram of wild type, mutant1- and mutant2-PIWIL1 protein. Both mutant1 and mutant2 are frameshift variants. **(C-D)** Immunostaining shows loss of PIWIL1 expression in MII oocytes (C) and 2-cell embryos (D) of *PIWIL1*$^{m1/m1}$. Nuclei were stained with DAPI. Scale bars = 50 µm.

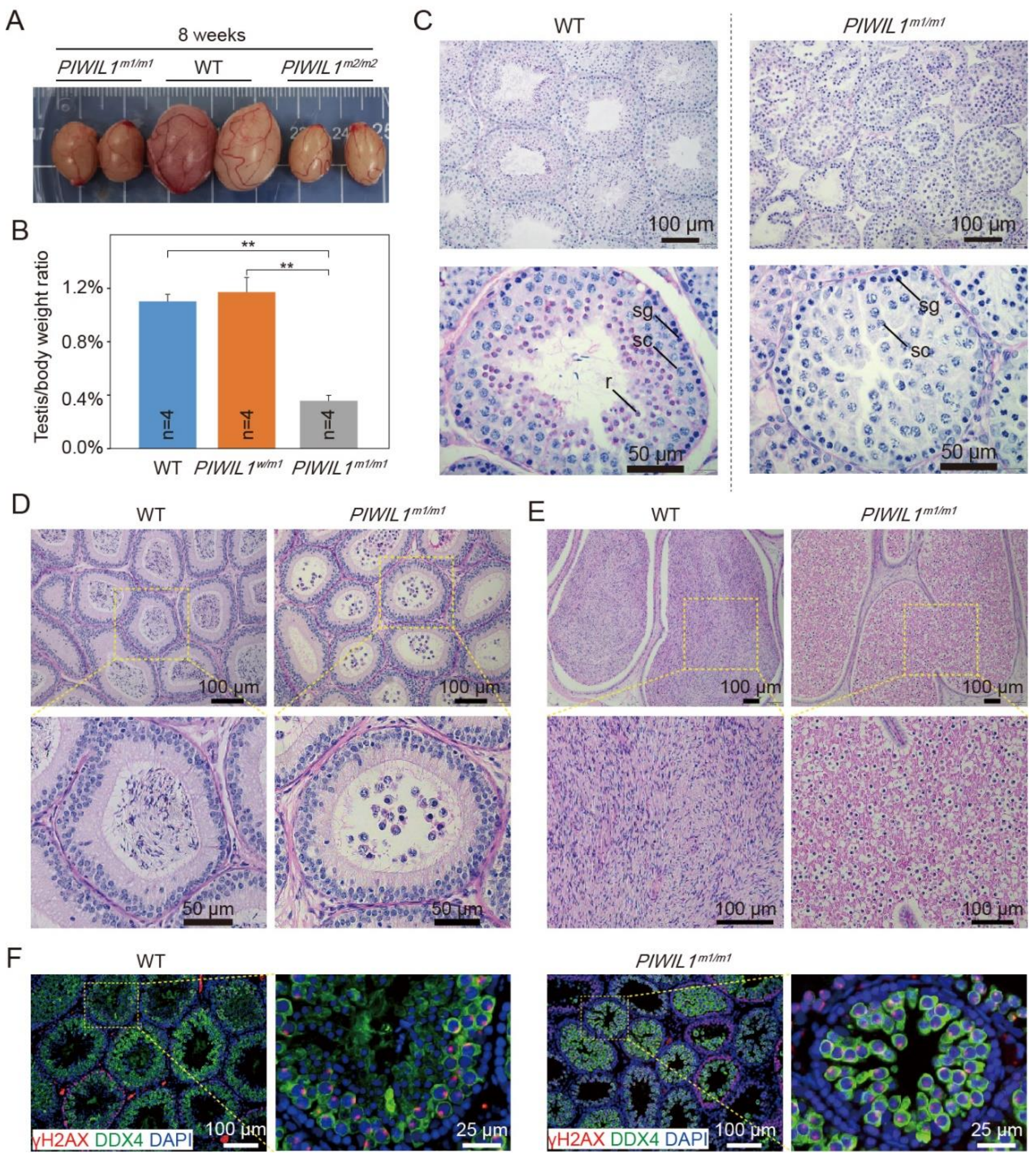


**Fig. s3 Spermatogenesis defects in *PIWIL1*-mutant golden hamsters**

**(A)** Comparison of the testes from 8-week-old wild-type, *PIWIL1*$^{m1/m1}$, and *PIWIL1*$^{m2/m2}$ golden hamsters. **(B)** The testis/body weight ratios. Testes are collected from 8-week-old wild-type, *PIWIL1*$^{w/m1}$ and *PIWIL1*$^{m1/m1}$ golden hamsters. n represents the number of hamsters analyzed. Error bars indicate s.e.m. ** P<0.05. **(C-E)** Periodic acid-Schiff (PAS) staining of adult testes (C), caput epididymis (D), and cauda epididymis (E). sg, spermatogonia; sc, spermatocyte; r, round spermatid. **(F)** Immunostaining of adult testes with DDX4 and γH2AX antibodies. Nuclei were stained with DAPI. Small γH2AX foci can be detected clearly in almost all spermatocytes in testes of the 5-week-old *PIWIL1*$^{m1/m1}$.

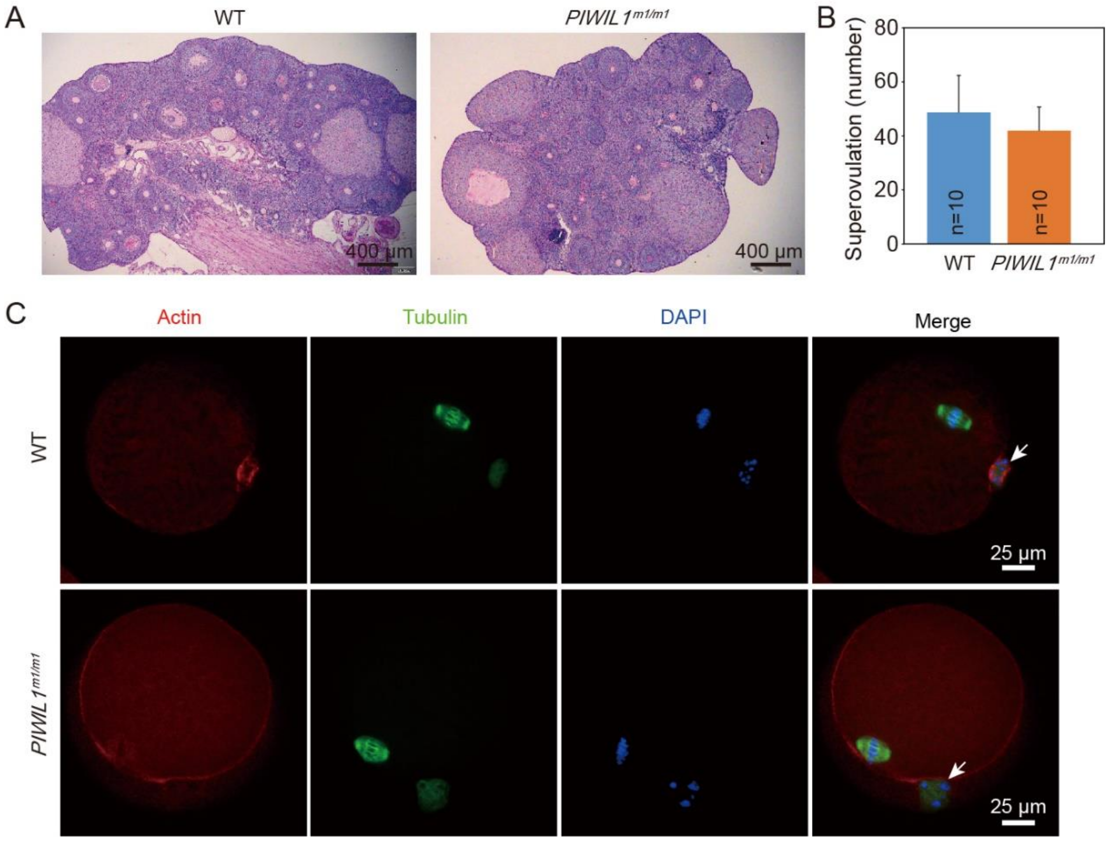


**Fig. s4 *PIWIL1* mutant shows no obvious phenotypic abnormality in follicle production or MII spindle formation**

**(A)** PAS staining of wild type and *PIWIL1*$^{m1/m1}$ ovarian sections. **(B)** The average numbers of ovulated oocytes collected from wild-type and *PIWIL1*$^{m1/m1}$ females, as determined by a superovulation assay. n is the number of superovulated golden hamsters. Error bars indicate s.e.m. **(C)** Immunofluorescence staining of the spindle in MII oocytes using ACTIN- and Tubulin-specific antibodies. Nuclei were stained with DAPI. Arrows indicate polar bodies.

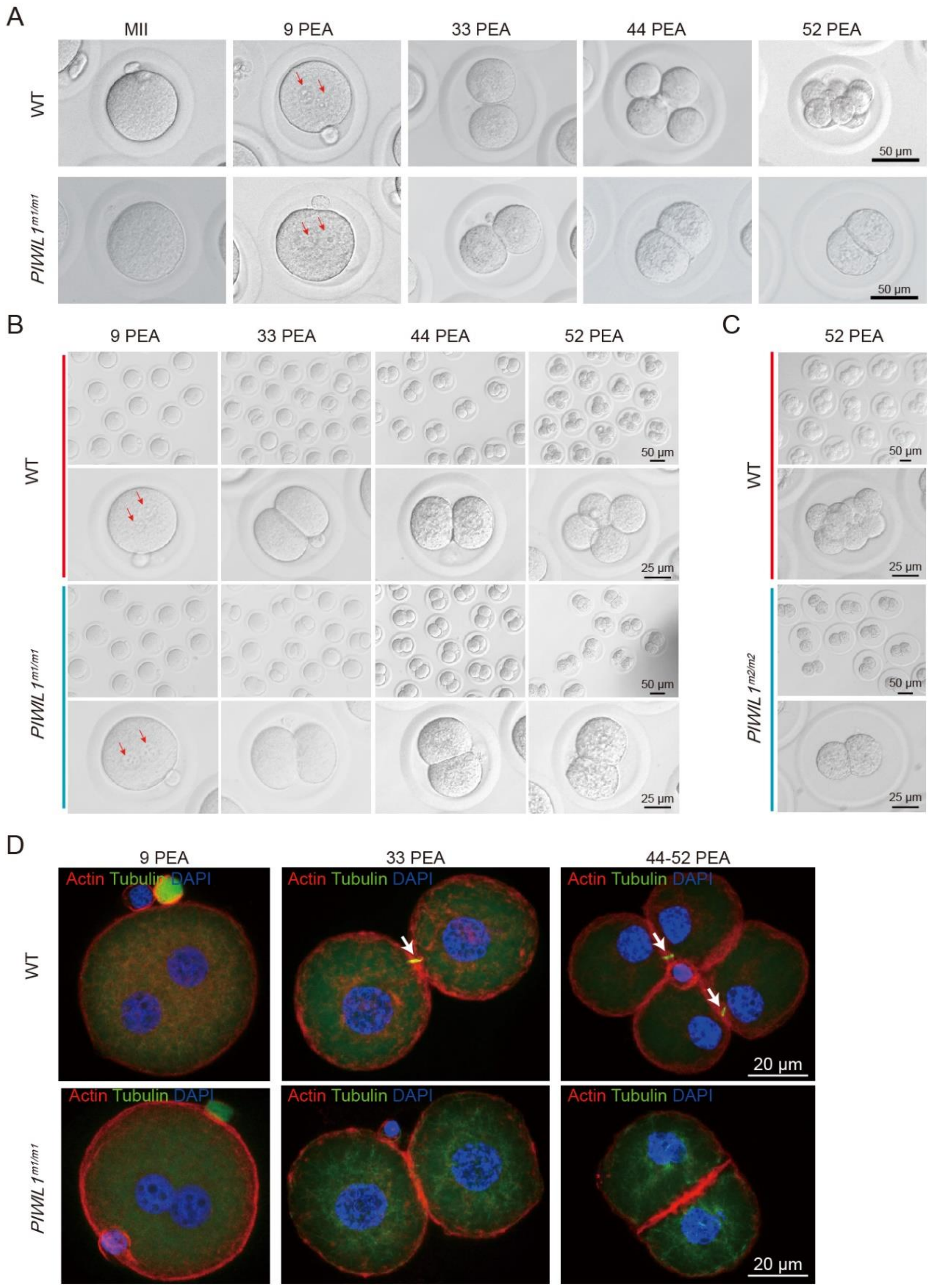


**Fig. s5 Development of *PIWIL1* mutant embryos was arrested at two-cell stage**

**(A)** Representative images of MII oocytes and embryos collected from the oviducts of wild-type and *PIWIL1*$^{m1/m1}$ females mated with wild-type males at 0, 9, 33, 44, and 52 PEA. Red arrows, pronuclei. **(B)** Representative images of *in vitro* cultured embryos obtained from wild-type and

*PIWIL1*$^{m1/m1}$ oocytes fertilized *in vivo* with wild-type sperm at 9, 33, 44, and 52 PEA. Red arrows, pronuclei. **(C)** Representative images of embryos collected from the oviducts of wild-type and *PIWIL1*$^{m2/m2}$ females at 52 PEA. **(D)** Immunofluorescence staining shows the absence of the microtubule bridge (white arrow) in embryos produced from *PIWIL1*$^{m1/m1}$ females mated with wild-type males.

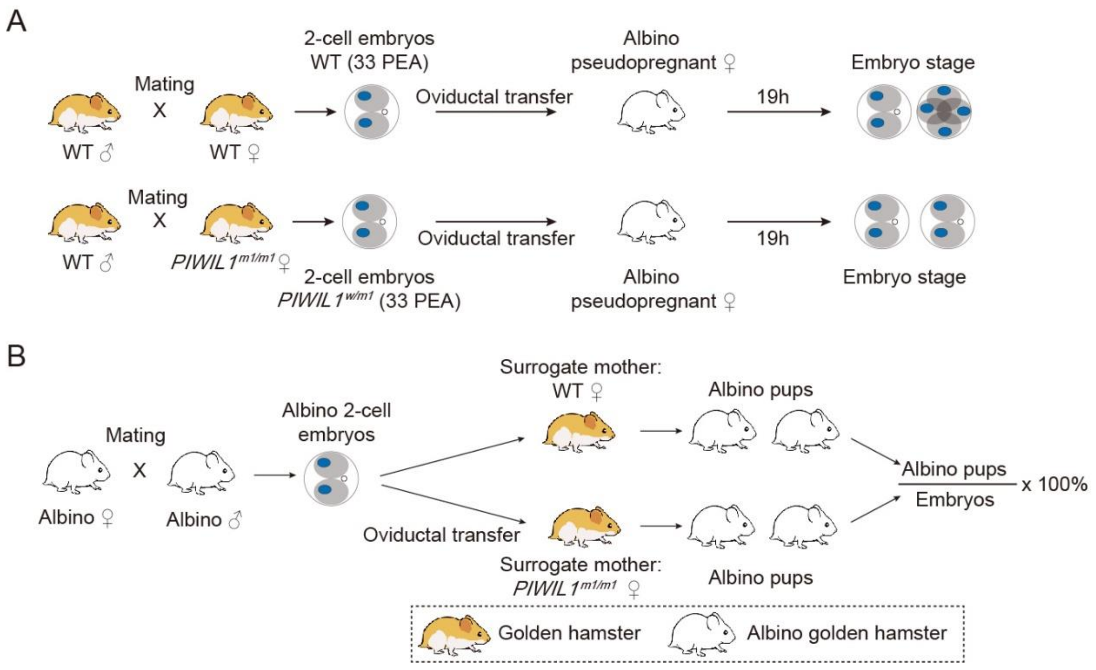


**Fig. s6 Strategies for oviductal transfer experiments**

**(A)** Wild-type or *PIWIL1*$^{m1/m1}$ two-cell embryos (33 PEA) were collected and transferred into the oviducts of pseudo-pregnant albino recipients. The transferred embryos were re-collected and examined at 52-54 PEA. **(B)** Albino wild-type two-cell embryos were collected and transferred into the oviducts of wild-type or *PIWIL1*$^{m1/m1}$ surrogate females, respectively.

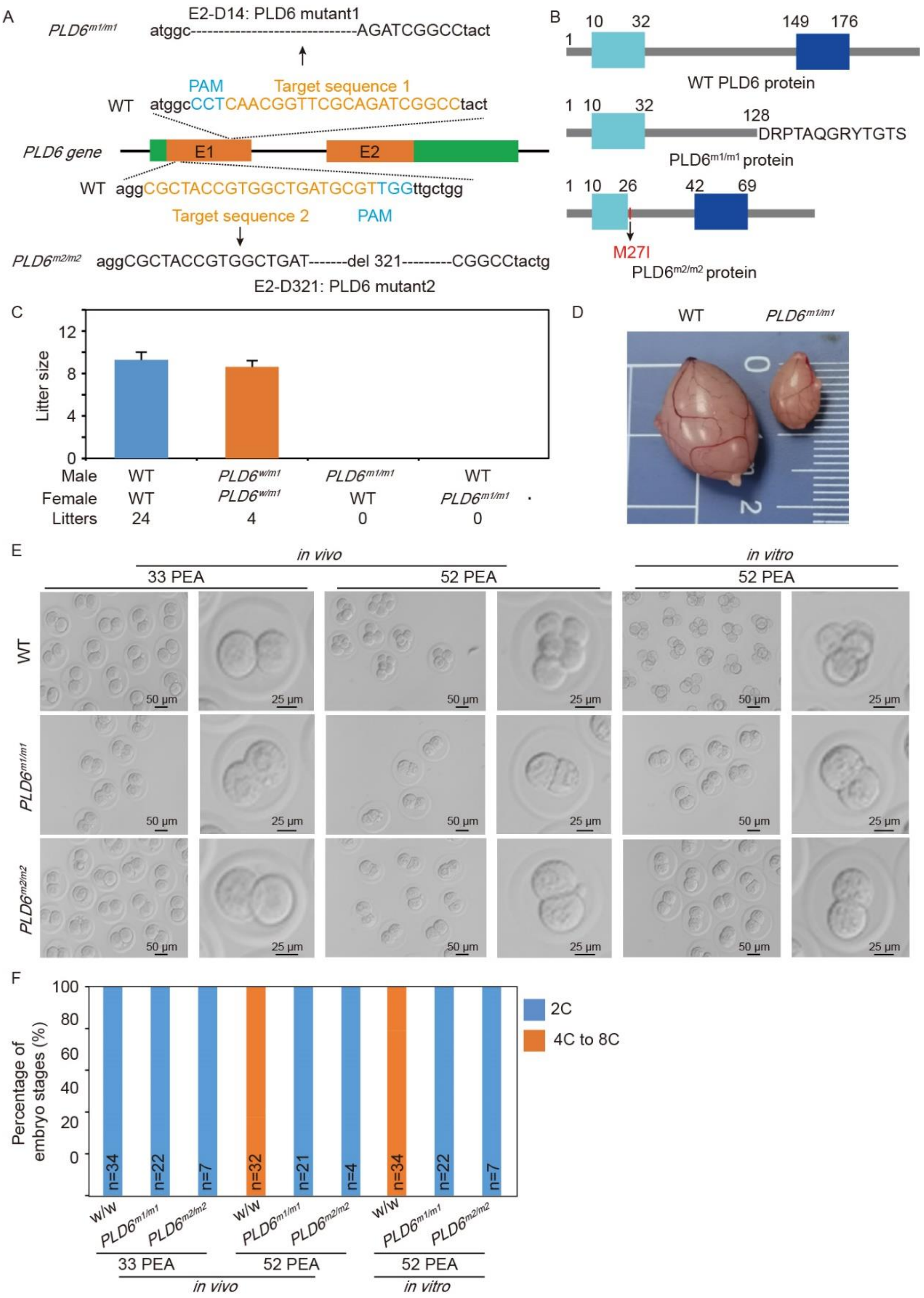


**Fig. s7 Embryos of *PLD6* mutant are arrested at the two-cell stage**

**(A)** Structure of golden hamster the *PLD6* gene and generation of *PLD6* mutants. The *PLD6* mutant1 (*PLD6*$^{m1/m1}$) and mutant2 (*PLD6*$^{m2/m2}$) contained 14 and 321 nt deletions in exon 1, respectively. **(B)** Diagram of wild-type and PLD6 mutant proteins. *PLD6*$^{m1/m1}$ caused a frameshift that generated a premature stop codon in *PLD6* mRNAs, while $PLD6^{m2/m2}$ caused a truncated PLD6 protein with a 107-amino acid deletion. **(C)** Fecundity of male and female $PLD6^{mut1(-/-)}$ golden hamsters. Error bars indicate s.e.m. **(D)** Comparison of the testes from 8-week-old wild-type and $PLD6^{m1/m1}$ golden hamsters. **(E)** Representative images of embryos produced *in vivo* and *in vitro*. $PLD6^{m1/m1}$ and $PLD6^{m2/m2}$ embryos were arrested at the two-cell stage. **(F)** Embryonic development of *PLD6* mutants was arrested at the 2-cell stage *in vivo* and *in vitro*. Embryos from wild-type and *PLD6* mutant females mated with wild-type males were collected at 33 or 52 PEA to determine the *in vivo* embryogenesis ratio. For *in vitro* analysis, zygotes from wild-type and *PLD6* mutant females mated with wild-type males were collected at 9 PEA and cultured *in vitro;* the embryogenesis ratio was determined at 33 and 52 PEA.

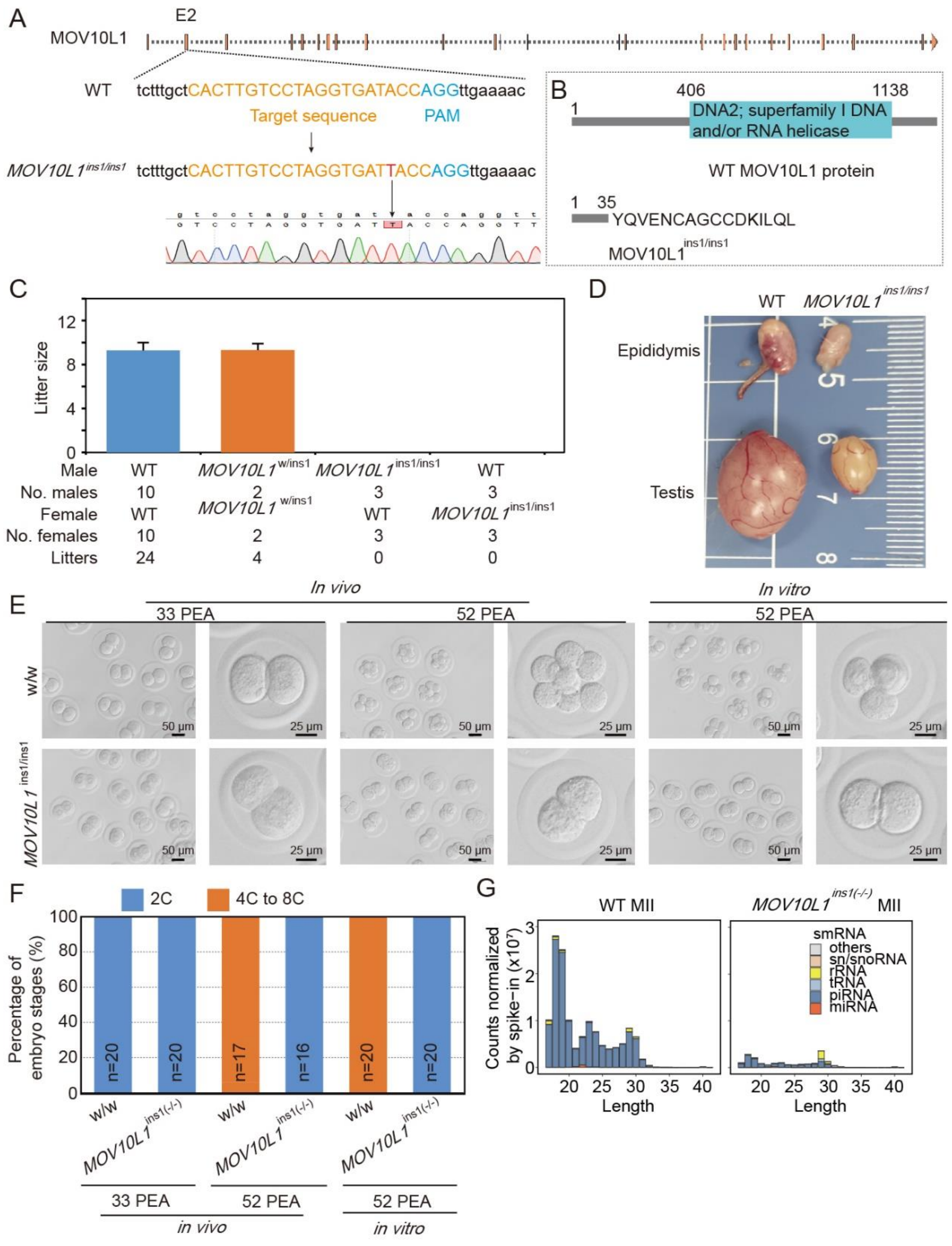


**Fig. s8 Embryos of *MOV10L1* mutant are arrested at the two-cell stage**

**(A)** Structure of golden hamster the *MOV10L1* gene and generation of *MOV10L1* mutants (*MOV10L1*$^{ins1/ins1}$). The *MOV10L1*$^{ins1/ins1}$ contained one thymine (T) insertion in exon 12. **(B)** Diagram of wild-type and MOV10L mutant protein. *MOV10L1*$^{ins1/ins1}$ caused a frameshift that generated a premature stop codon in *MOV10L1* mRNAs. **(C)** Fecundity of male and female *MOV10L1*$^{ins1/ins1}$ golden hamsters. Error bars indicate s.e.m. **(D)** Comparison of the testes from 8-

week-old wild-type and *MOV10L1*$^{ins1/ins1}$ golden hamsters. **(E)** Representative images of embryos produced *in vivo* and *in vitro*. *MOV10L1*$^{ins1/ins1}$ embryos were arrested at the two-cell stage. **(F)** Embryonic development of *MOV10L1*$^{ins1/ins1}$ was arrested at the two-cell stage *in vivo* and *in vitro*. Embryos from wild-type and *MOV10L1*$^{ins1/ins1}$ females mated with wild-type males were collected at 33 or 52 PEA and the *in vivo* embryogenesis ratio was determined. For *in vitro* analysis, zygotes from wild-type and *MOV10L1*$^{ins1/ins1}$ females mated with wild-type males were collected at 9 PEA and cultured *in vitro*; the embryogenesis ratio was determined at 33 and 52 PEA. **(G)** Composition of small RNAs in wild-type and *MOV10L1*$^{ins1(-/-)}$ MII oocytes.

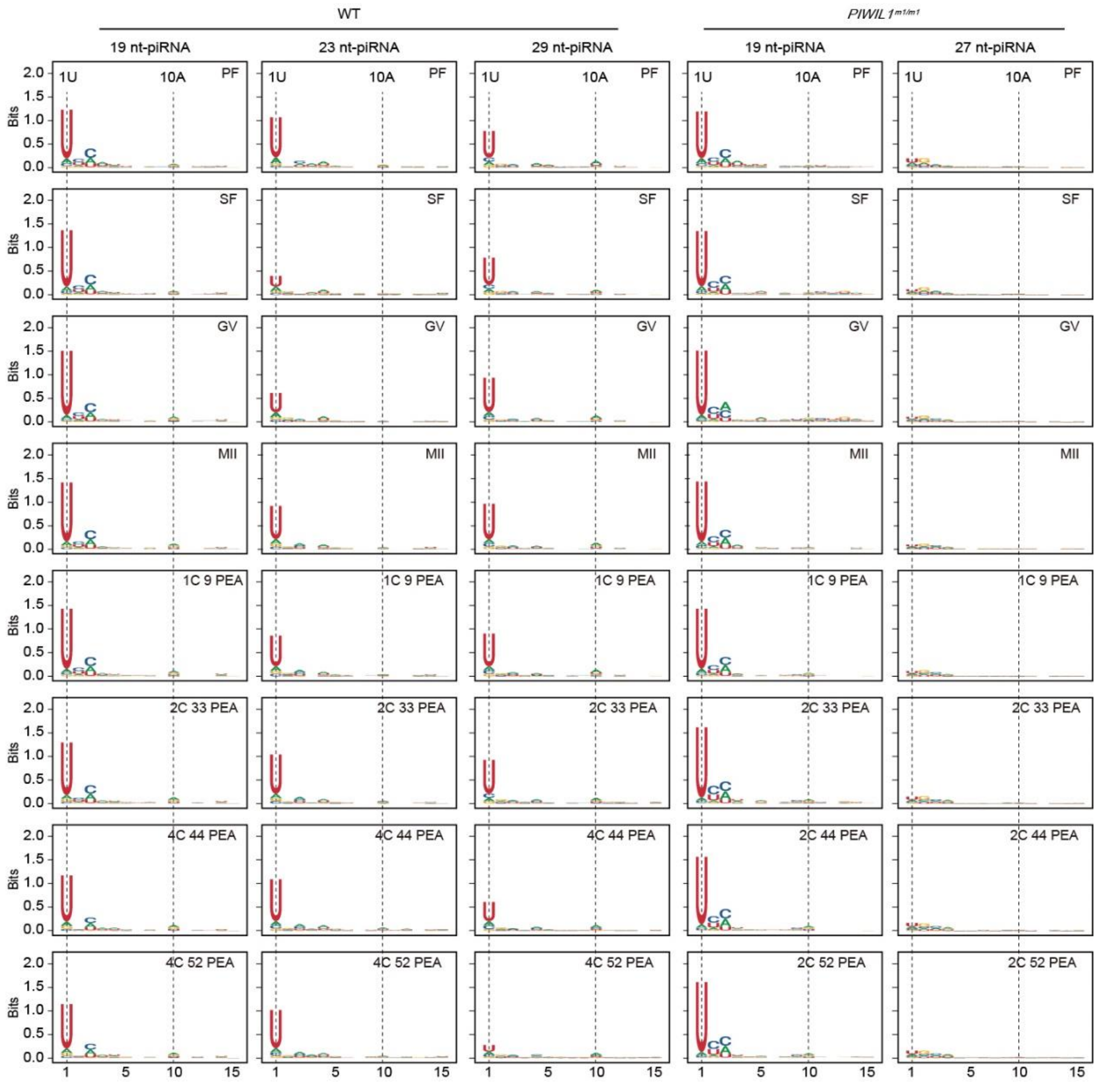


**Fig. s9 Sequence analysis of piRNAs expressed in oocytes and embryos of wild-type and *PIWIL1*-mutant females**

The piRNAs identified in oocytes and embryos collected from wild-type females showed three main peaks in their size distribution: 18-20 nt (19 nt-piRNA), 22-24 nt (23 nt-piRNA), and 28-30 nt (29 nt-piRNA). All three piRNA populations preferentially carried a 5′ uracil (U). Oocytes and embryos collected from *PIWIL1*$^{m1/m1}$ females only harbored two piRNA populations. The piRNAs of 18-20 nt showed a strong preference for a 5′ U, while 26-28 nt piRNAs (27 nt-piRNA; expressed at low levels) showed a weak preference for a 5′ U.

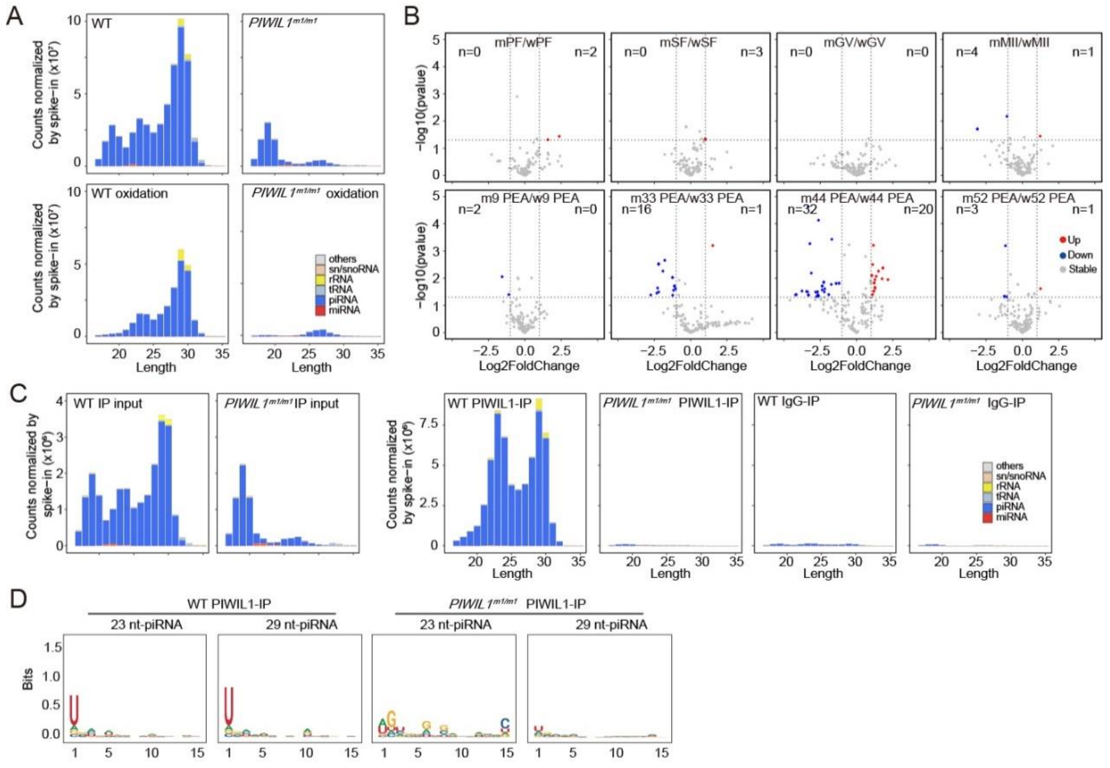


**Fig. s10 Two populations of piRNAs with peaks at 23-nt and 29-nt are associated with PIWIL1**

**(A)** Composition of small RNAs in wild-type and *PIWIL1*$^{m1/m1}$ MII oocytes with or without NaIO4 treatment. **(B)** Differential analysis of miRNA expression during oogenesis and early embryo development. miRDeep2 was used for *de novo* miRNA identification. m, *PIWIL1* mutant; w, wild-type. **(C)** Size distribution of PIWIL1-associated piRNAs identified by immunoprecipitation with a PIWIL1-specific antibody in wild-type and *PIWIL1*$^{m1/m1}$ MII oocytes. Rabbit non-specific immunoglobulin G (IgG) antibody served as a negative control. Two populations of piRNAs, 22-24 nt and 28-30 nt, bound to PIWIL1. The small RNA counts in (A) and (C) were normalized by exogenous spike-in. **(D)** Sequence analysis of PIWIL1-piRNA in MII oocytes.

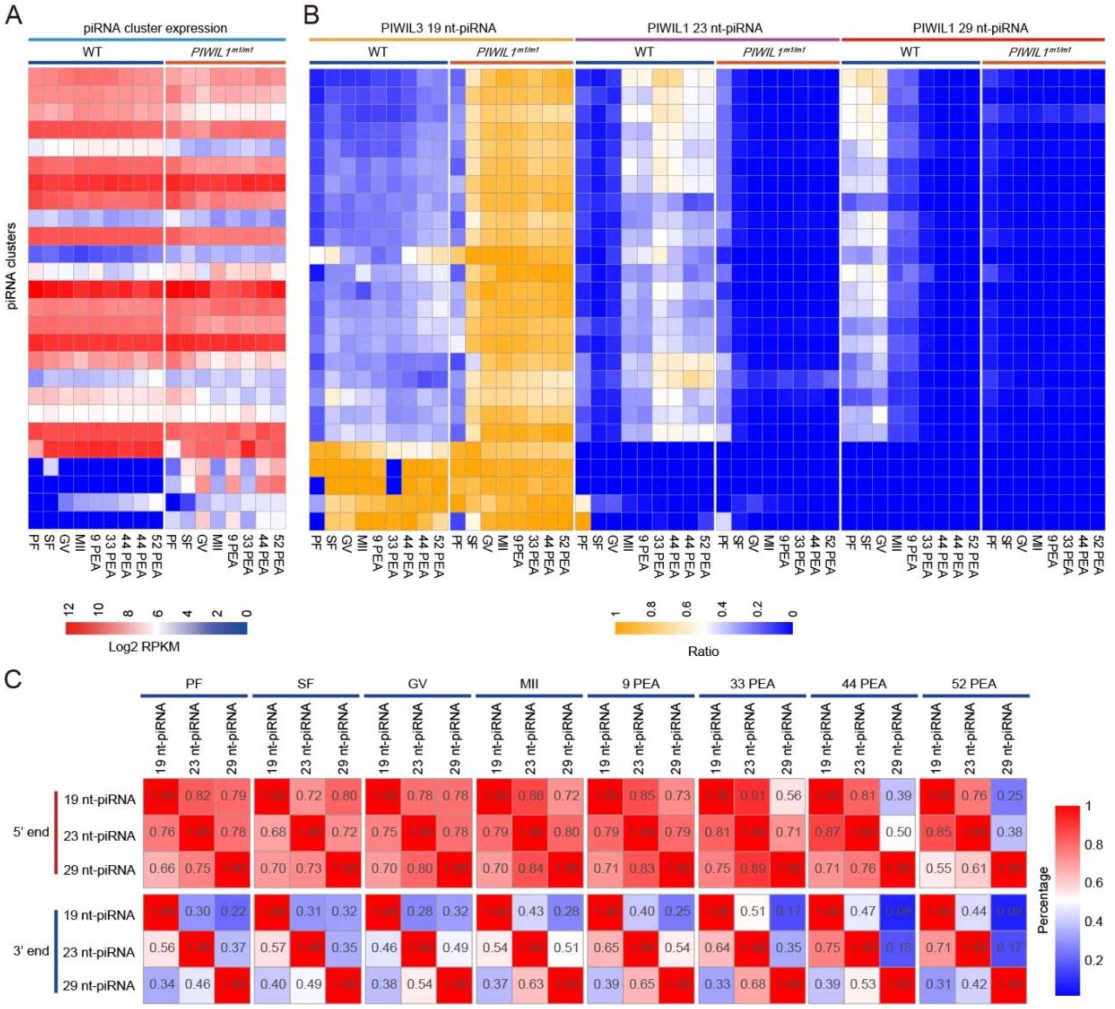


**Fig. s11 PIWIL1-associated 23 nt- and 29 nt-piRNAs are co-localized in the genome.**

**(A)** Heatmaps show the expression level (RPKM) of top piRNA clusters, which generated >90% of the unique mapped piRNAs in each oocyte and embryo at different developmental stages. **(B)** The relative abundances of PIWIL1 23nt-, PIWIL1 29nt- and PIWIL3 19nt-piRNAs among total piRNAs (A) in each cluster are shown. Almost all of the piRNA clusters that produce PIWIL1 23 nt- and 29-nt piRNAs are identical, while several piRNA clusters uniquely produced PIWIL3 19 nt-piRNAs. **(C)** The ratio of piRNAs with identical 5' ends (up) and 3' ends (bottom) among PIWIL1 23nt-, PIWIL1 29nt- and PIWIL3 19nt-piRNAs. Only uniquely mapped piRNAs are calculated in (A) and (B).

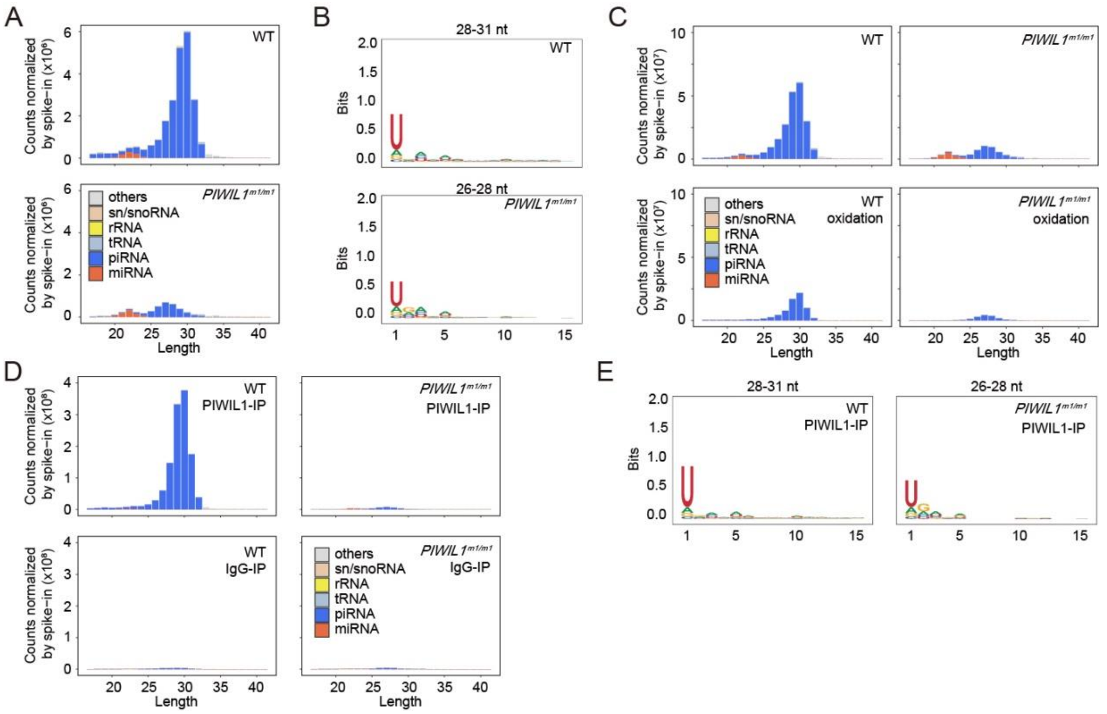


**Fig. s12 Characteristics of small RNAs in the testes of wild-type and *PIWIL1*$^{m1/m1}$ males**

**(A)** Composition of small RNAs in the testes of 3-month-old wild-type and *PIWIL1*$^{m1/m1}$ males. Small RNA reads were classified into tRNAs, rRNAs, sn/snoRNAs, miRNAs, and piRNAs, as indicated by different colors. **(B)** Sequence analysis of piRNAs expressed in testes. The 28-31 nt piRNAs found in wild-type testes and the remaining 26-28 nt piRNAs in *PIWIL1*$^{m1/m1}$ testes both showed a strong preference for 5′ U. **(C)** Composition of small RNAs in wild-type and *PIWIL1*$^{m1/m1}$ testes with or without $NaIO_4$ oxidation treatment. **(D)** Size distribution of PIWIL1-associated piRNAs in wild-type and *PIWIL1*$^{m1/m1}$ testes immunoprecipitated with PIWIL1-specific antibody. Rabbit non-specific immunoglobulin G (IgG) served as a negative control. **(E)** Sequence analysis of PIWIL1- piRNAs expressed in MII oocytes. Both the PIWIL1 29 nt- and PIWIL1 23 nt-piRNAs showed a strong preference for 5′ U. The small RNA counts were normalized by exogenous spike-in in (A), (C), and (D).

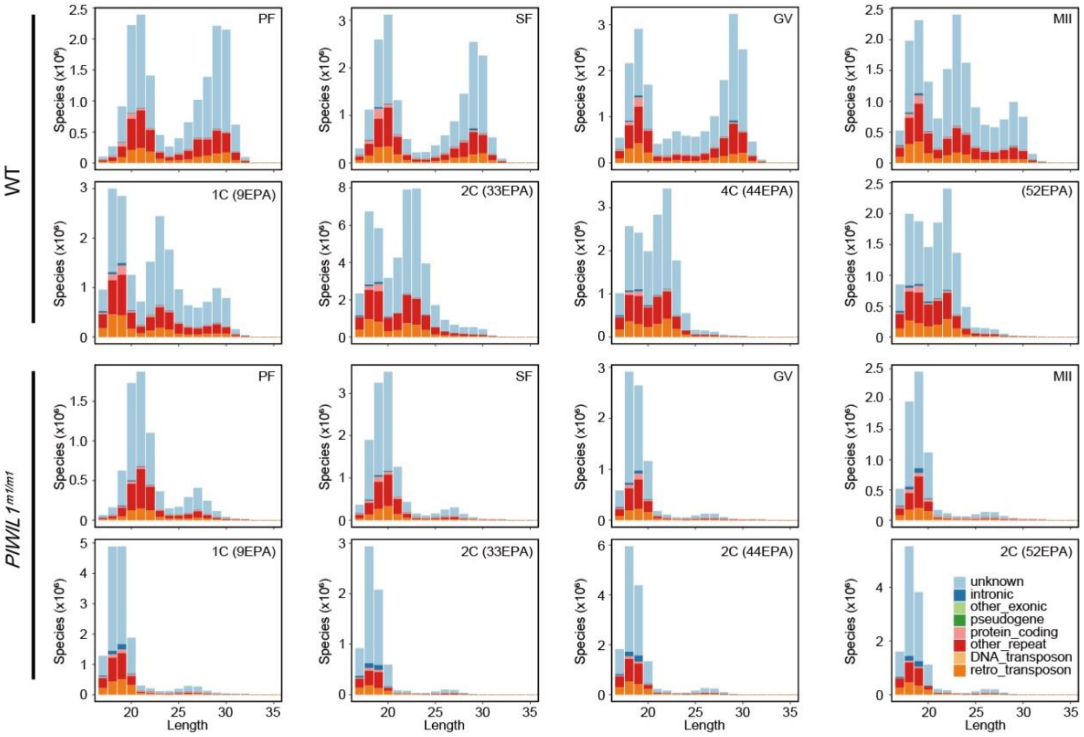


**Fig. s13 Genomic annotation of piRNAs at different stages**

Genomic annotation of piRNA species of different sizes identified in oocytes at the primary follicle stage through embryos at 52 PEA in wild-type and *PIWIL1*$^{m1/m1}$ golden hamsters.

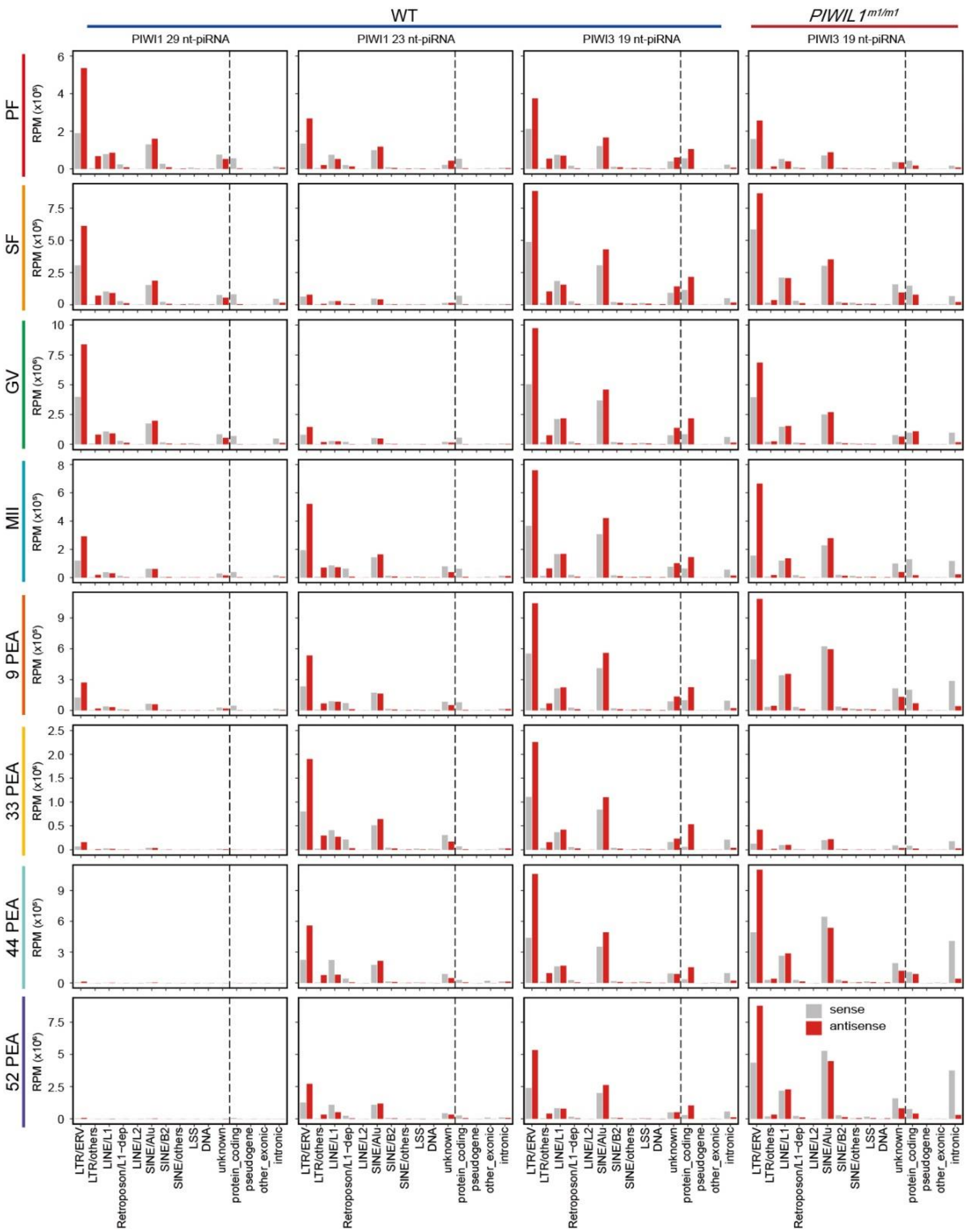


**Fig. s14 Distribution of piRNAs generated from different genomic regions**

Bar graphs show the distribution of piRNAs generated from different genomic regions. PIWIL1 29 nt-piRNAs, PIWIL1 23 nt-piRNA, and PIWIL3 19 nt-piRNAs were analyzed separately. Histograms left of the vertical dashed line show different families of repeat elements; histograms right of the vertical dashed line show the gene-related regions. The combination of Low complexity, Simple_repeat, and Satellite is designated as LSS. In *PIWIL1* $^{m1/m1}$, only PIWIL3 19 nt-piRNAs are shown.

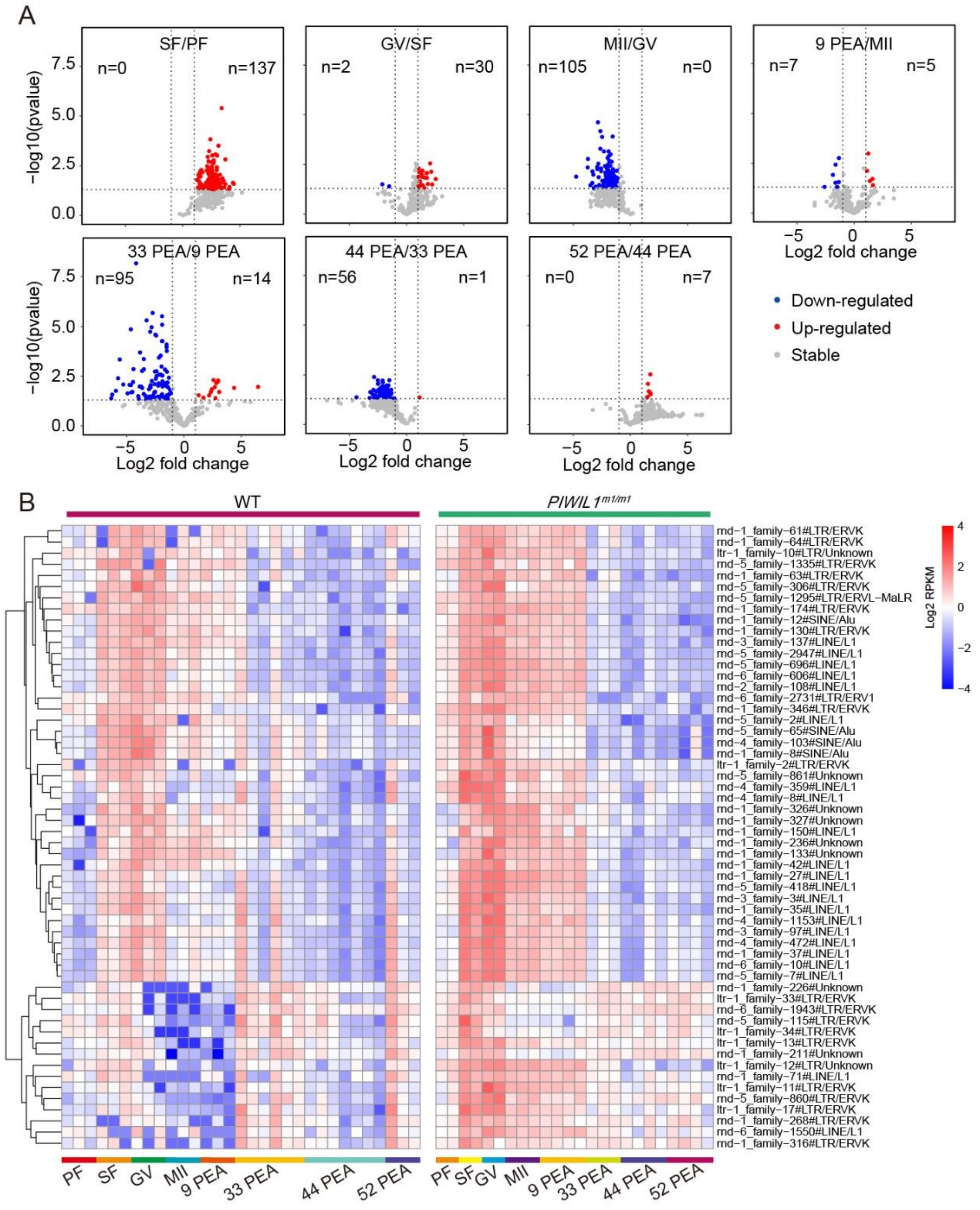


**Fig. s15 TE expression levels during oogenesis and embryo development**

**(A)** Volcano plots show the differentially expressed TEs between two adjacent stages during oogenesis and embryo development in wild-type golden hamsters. The highly significant up- or down-regulated TEs (≥ 2 folds; Welch two sample *t*-test, p-value < 0.05) are indicated in red or blue, respectively, with TE numbers shown at the top. **(B)** Heatmap of up-regulated TE expression levels in wild-type and *PIWIL1*$^{m1/m1}$ oocytes and embryos at different stages. The highly significant up-regulated TEs (≥ 2 folds) in *PIWIL1*$^{m1/m1}$ compared to wild-type at each stage from oocytes at PF to embryos at 9 PEA are plotted.

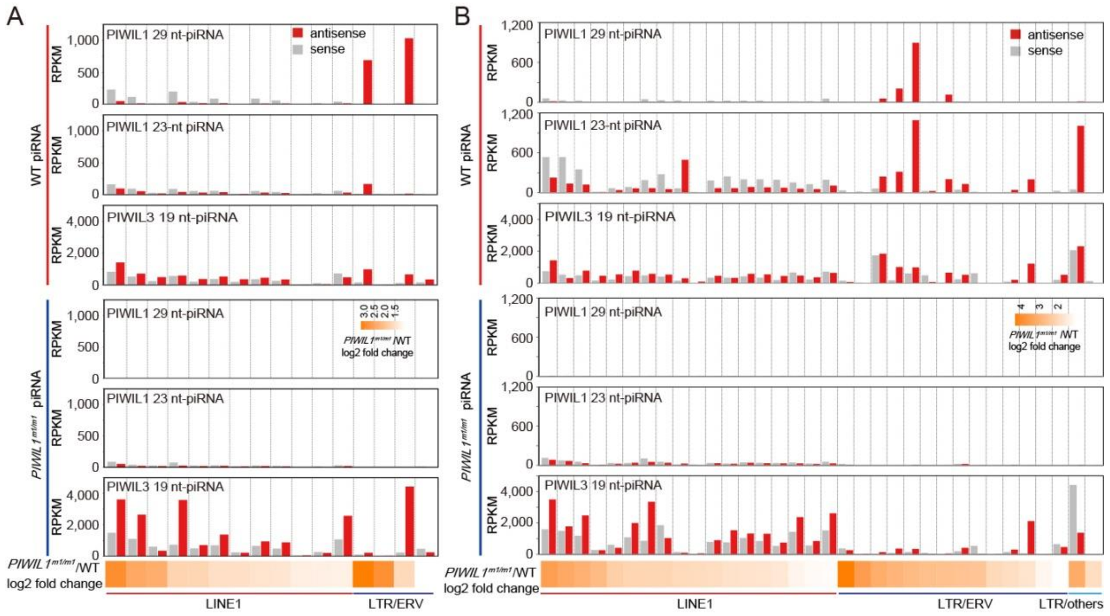


**Fig. s16 Anti-sense piRNAs were enriched with up-regulated TEs**

**(A-B)** The bar graph shows the expression level (RPKM) of piRNAs mapped to the sense (gray) or anti-sense (red) directions of each TE family in GV oocytes (A) and 1-cell embryos at 9 PEA (B). Different TE family-derived piRNAs are plotted. The significantly up-regulated TE families are listed with log2 fold change in expression level between *PIWIL1* mutant versus wild-type MII oocytes. The fold change level is indicated by differences in orange hue in the heatmap.

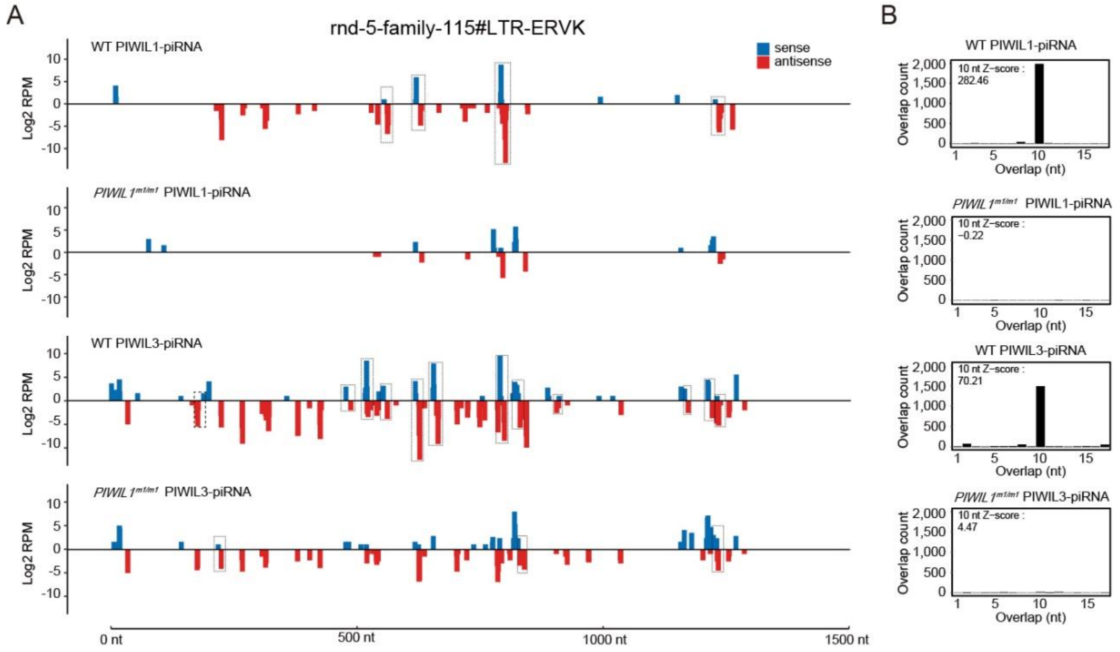


**Fig. s17 Representative examples of piRNAs derived from up-regulated ERV**

**(A)** Examples of PIWIL1-piRNA and PIWIL3-piRNA distribution in ERV families that were up-regulated in *PIWIL1*$^{m1/m1}$ MII oocytes. The expression levels (RPKM) represent the normalized number of all mapped piRNAs with the same 5' end at each position. The dotted boxes indicate the Ping-Pong signal between the adjacent sense and antisense piRNAs. **(B)** The 5'-5' overlap between piRNA sense- and antisense strands was analysed to determine the presence of Ping-Pong signature. The number of pair of piRNA reads at each position is plotted. Significance of 10-nt overlap ('Ping-Pong') was determined from *Z*-score. *Z*-score >1.96 corresponds to p-value < 0.05.

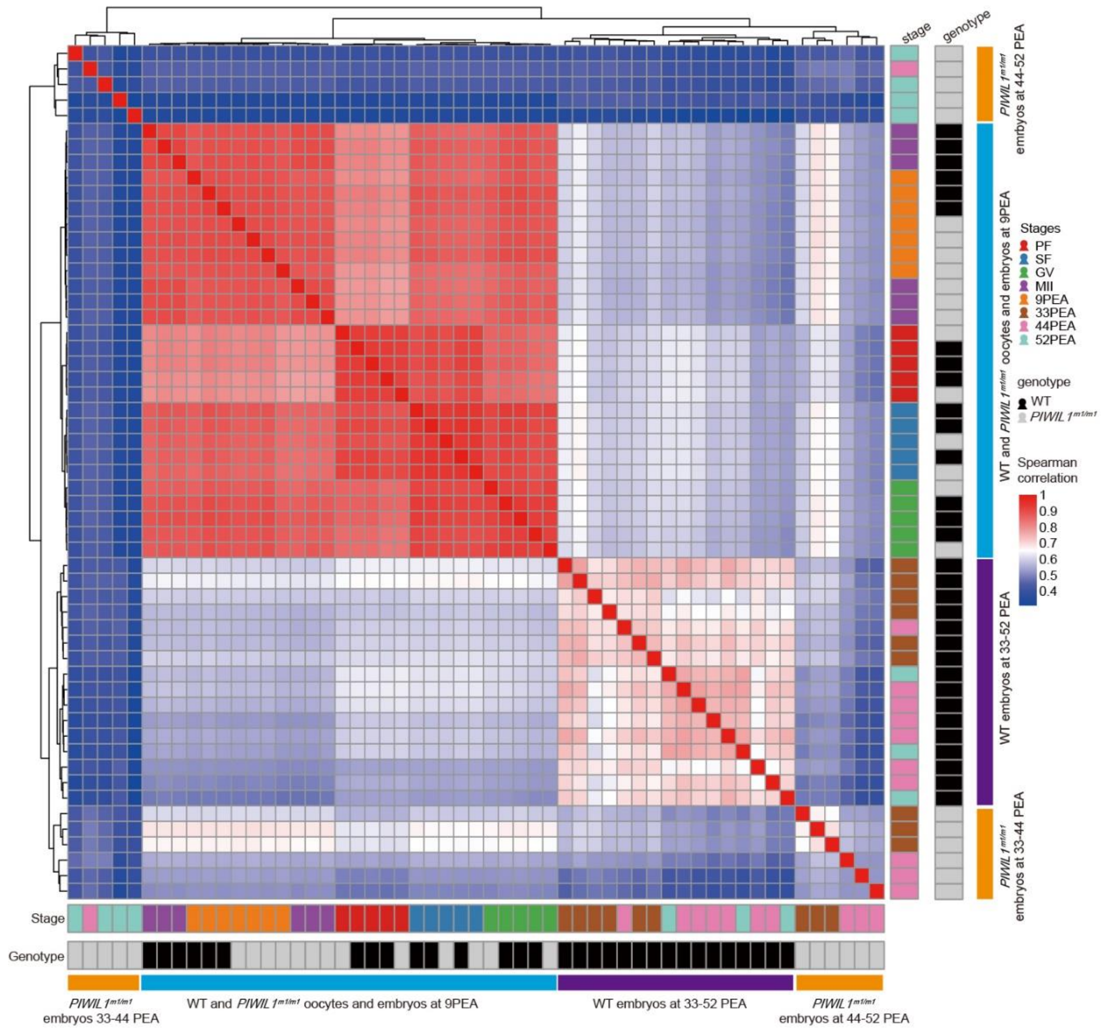


**Fig. s18 Correlation of gene expression between oocytes and embryos at different stages**

Heatmap of Spearman correlation coefficient (R) of gene expression in oocytes and embryos at different stages collected from wild-type and *PIWIL1*$^{m1/m1}$ golden hamsters.

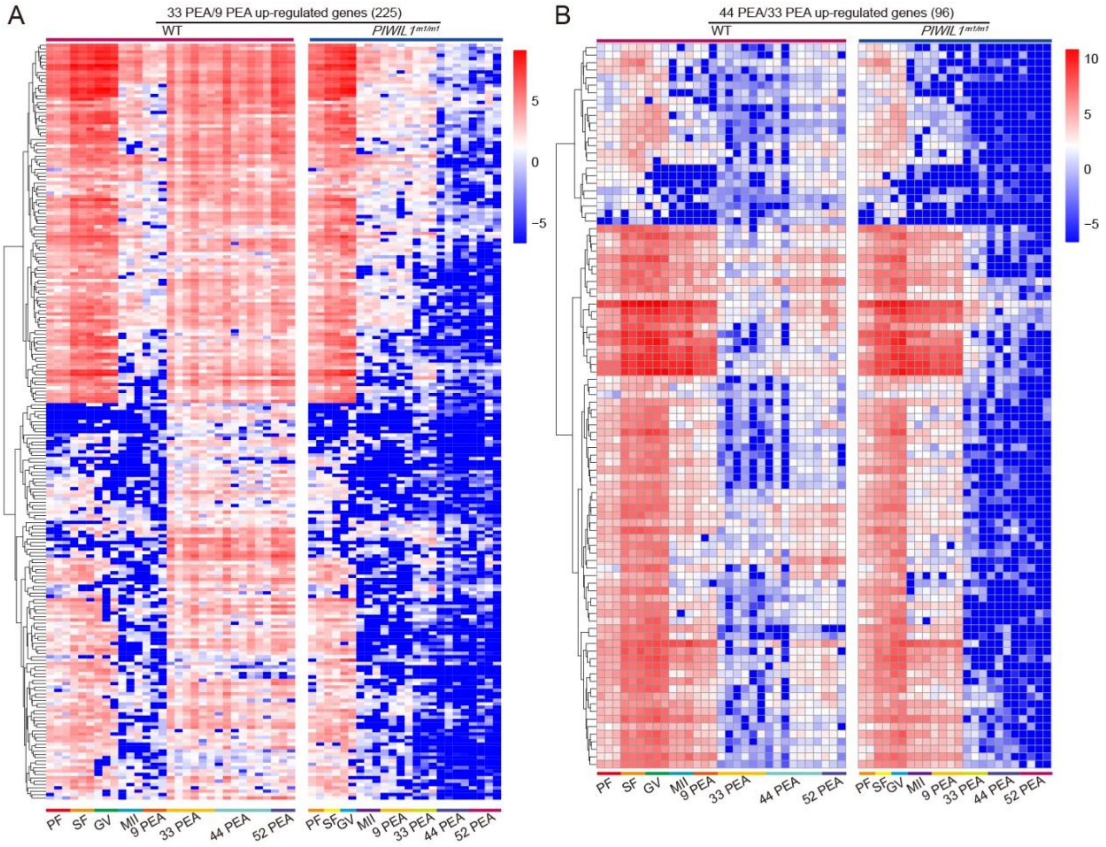


**Fig. s19 ZGA is impaired in *PIWIL1*-mutant embryos**

**(A-B)** Heatmap of gene expression at different stages of wild-type and *PIWIL1*-mutant oocytes and embryos. Only the genes which were up-regulated from 9 PEA to 33 PEA (A) or 33 PEA to 44 PEA (B) stages in wild-type embryos are shown. The up-regulation of gene expression in early embryogenesis of wild-type embryos was barely detectable in *PIWIL1* mutants.

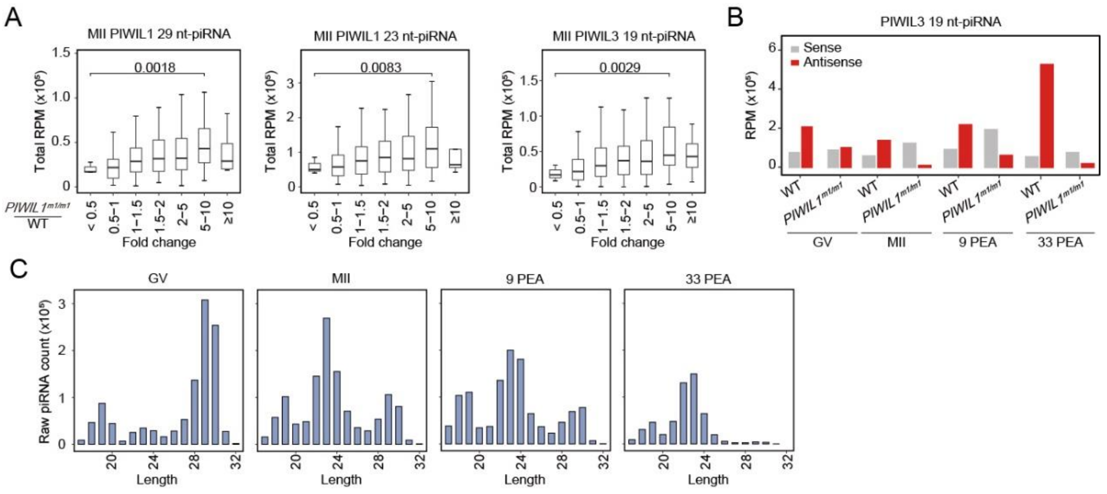


**Fig. s20 PIWIL1-piRNAs participate in the degradation of maternal mRNAs**

**(A)** The expression levels of PIWIL1- and PIWIL3-piRNAs were correlated with the up-regulation levels of their target mRNAs. The method used to calculate the level of gene up-regulation is the same as in Figure 4E. **(B)** The size distribution of piRNAs down-regulated in *PIWIL1*$^{m1/m1}$ compared to wild-type. **(C)** Bar graph shows the count of PIWIL3 19 nt-piRNAs derived from sense and anti-sense strands of mRNAs in *PIWIL1*$^{m1/m1}$ and wild-type oocytes and embryos at different stages.
